# Mechanical and chemical macrophage activation differ by arm, not by amplitude

**DOI:** 10.64898/2026.11.10.765627

**Authors:** Chunlian Zhong, Weiya Chen, Tao Zeng, Shuying Zhang, Joey Zhu, Zhaoqing Cong

## Abstract

Mechanical cues are widely reported to polarize macrophages toward an M1-like state, but the claim rests on nuclear factor kappa B/activator protein 1 (NF-κB/AP-1) effector markers read without a common scale, and the literature does not agree on the sign of the stiffness effect. We asked a quantitative question instead: by what fraction of the chemical M1 response does mechanical perturbation move the interferon-stimulated gene factor 3 (ISGF3)-dependent interferon-stimulated gene module? Scoring ten independent studies of substrate stiffness and applied force against lipopolysaccharide (LPS) + interferon-gamma (IFN-γ) anchors, processed through an identical pipeline, we find a pooled shift of −0.9% of the chemical response (95% confidence interval −6.2% to +4.5%), equivalent to zero at a ±10% margin (p = 0.0019) with a minimum detectable effect of 8.8%. The result is unchanged across three independent ISG definitions. Nor is it a sensitivity failure: the most transcriptionally responsive mechanical contrast rewires thousands of genes while leaving the module inside a size-matched random distribution. Extending to ten further mechanical modalities shows mechanics is not inert toward these genes; shifts run in both directions, but no contrast supported by within-study inference exceeds 15% of the chemical anchor. Rescoring the same contrasts against an NF-κB effector module anchored on human LPS kinetics separates the two arms: across 21 modality contrasts the effector arm sits 17.7 percentage points above the interferon arm (paired p = 0.032), and in the four designs whose permutation space can reach p < 0.05 the separation is 34 points. Compression moves the effector arm +36% of the LPS anchor (p = 0.002) and the interferon arm −9% (p = 0.004). The separation is absent under static stiffness, where neither arm moves, and its sign is not fixed. A three-arm in-vivo magnetomechanical dataset assigns the effect to the rotating field rather than the internalized particle, raising the effector arm while lowering the interferon arm. Mechanically induced “M1-like” states are not the chemical M1 program rescaled. Where mechanical loading registers at all, it moves the effector arm and not the interferon arm: interferon-stimulated genes are the wrong readout for mechanical perturbation of macrophages.

## 1 Introduction

Macrophage activation states are usually named by reference to two chemical protocols. “M1” denotes the state reached by LPS together with IFN-γ, and at the transcriptional level that state has two distinguishable arms: an NF-κB/AP-1 effector arm carrying inducible nitric oxide synthase 2 (NOS2), tumor necrosis factor-alpha (TNF-α), interleukin 6 (IL6), and interleukin 12 subunit beta (IL12B), and an interferon arm carrying the ISGF3-dependent interferon-stimulated genes. The canonical stimulus induces both arms together. Because they arrive together, they are rarely distinguished. Marker panels assembled for convenience (iNOS, TNF-α, IL-6, CD80 by qPCR, flow cytometry or ELISA) sample the effector arm almost exclusively. A state that scores positive on such a panel is routinely called M1 without the interferon arm ever being examined, a problem already visible in transcriptome-based spectrum models of human macrophage activation (1).

Over the past decade macrophages have also been shown to respond to purely mechanical inputs. Substrate stiffness, surface topography, three-dimensional confinement, cyclic stretch, fluid shear, hydrostatic pressure, ultrasound and matrix viscoelasticity all alter macrophage gene expression. Candidate transducers have been identified. Piezo-type mechanosensitive ion channel component 1 (PIEZO1) couples mechanical load to inflammatory output in myeloid cells (2,3), transient receptor potential cation channel subfamily V member 4 (TRPV4) has been implicated in stiffness-dependent activation (4), and Yes-associated protein/Transcriptional coactivator with PDZ-binding motif (YAP/TAZ), the canonical stiffness-responsive transcriptional coactivators, restrain TANK-binding kinase 1 (TBK1) and so sit plausibly upstream of interferon output (5). A large and growing literature summarizes these observations with the statement that mechanical cues polarize macrophages toward M1.

That statement is made on the effector-arm panels described above, and it is made without a common scale. That absence isn’t a presentational shortcoming: it is a measurement problem, and it is the reason the field’s disagreements have proved impossible to settle. When one laboratory reports that a stiff substrate “induces M1” on the basis of a two-fold rise in NOS2 transcript and another reports that a soft substrate does the same on the basis of a 40% rise in secreted TNF-α, the two claims are not commensurable. They are expressed in different units, read from different arms of the program, and normalized to different baselines. Nothing in either report tells a reader whether the state reached resembles the state produced by LPS plus IFN-γ by 5% or by 95%. The quantity that would settle the question, the size of the mechanical effect expressed in units of the chemical effect, measured the same way, is almost never reported. Almost no mechanical study processes a chemical arm in the same batch.

Three consequences follow. First, the literature does not agree on sign: stiff substrates have been reported to drive M1 (4,6), soft substrates to drive M1 (7), and stiffness has also been reported to push tumor-associated macrophages toward M2 (8). Second, almost no mechanical study processes a chemical reference arm alongside the mechanical arm. A reported “M1-like” shift therefore cannot be placed on any scale: a reader cannot distinguish a shift that is 5% of the chemical response from one that is comparable to it. Third, a negative result in this field is currently unpublishable in any useful form. “We did not detect induction” carries no bound.

The interferon arm offers a way out of this, for a reason that is specific rather than convenient. Of the two arms of chemical M1, the interferon arm is the one most tightly tied to the defining stimulus: it is driven by a well-characterized transcription factor complex, its target set has been mapped by chromatin occupancy rather than by expression correlation (9), and it is large enough to be scored robustly in a single sample. That same work shows that IFN-γ also recruits ISGF3 extensively, so the relevant set is the ISGF3-dependent one rather than a generic “type I interferon” list. Otherwise an LPS + IFN-γ reference arm would be partly self-referential. If mechanically induced states engage the same program as chemical M1, the ISGF3-dependent module should move. If they engage only the effector arm, it should not.

There is also a practical argument for choosing this arm, distinct from the mechanistic one. An absence is only informative if it is bounded, and an absence can only be bounded on a scale whose unit is established independently of the experiment that fails to move it. The interferon arm supplies such a unit: the chemical stimulus that defines M1 moves the ISGF3 module by a large, repeatedly measurable amount in every cell source and on every platform we examined, so the denominator of the comparison is not a modelling assumption but a measurement. An NF-κB/AP-1 effector panel read by qPCR, by contrast, cannot supply a stable denominator. Its dynamic range depends on the reference gene, the time point and the assay, which is precisely why a decade of effector-arm reports has not converged. Choosing the interferon arm trades breadth of phenotypic coverage for the ability to state a bound. That bound is what the field currently lacks.

We asked a quantitative question instead of a qualitative one: by what fraction of the chemical M1 response does mechanical perturbation move the ISGF3-dependent module? Posing it this way converts an unfalsifiable absence into a bounded estimate. It also dictates the design. Every mechanical contrast is scored with a rank-based single-sample statistic that needs no cross-study normalization, and every mechanical shift is reported in the same units as a chemical anchor, LPS + IFN-γ or IFN-γ, scored by the identical pipeline. The anchor is not background information. It is the ruler, and we test its reproducibility before using it.

We assembled ten independent studies of substrate stiffness and directly applied force, together with 21 further datasets spanning ten additional mechanical modalities, and scored all of them against three independent ISG definitions and a YAP/TAZ signature. We report the pooled mechanical shift as a percentage of the chemical anchor with its confidence interval, establish the margin at which it is statistically equivalent to zero, and state the resolution the available evidence actually has. We then ask three questions: whether the same conclusion holds across mechanical modalities, whether the YAP/TAZ route can be adjudicated with current tools, and whether the placement of a magnetic actuator (on the plasma membrane or inside the lysosome) changes the answer.

Finally, we checked whether this comparison has been made before. In the OpenAlex citation graph, 845 works cite one of the two PIEZO1 macrophage-mechanics papers and 437 cite either the ISGF3 target definition or the YAP/TAZ–TBK1 study; three works cite both, and the only one relevant to macrophage activation is a review (10). Of the 845 mechanics-citing works, one carries a title suggesting cross-study synthesis, and it concerns idiopathic pulmonary fibrosis rather than macrophage polarization. Nine targeted searches return macrophage transcriptome meta-analyses that are chemical throughout: the macIDR compendium aggregates eight in-vitro cytokine-differentiated models (11), a 2024 consensus exercise addresses murine polarization signatures under cytokine stimulation (12), and a 2023 proteomic survey covers the MØ/M1/M2a/M2c continuum (13). None includes a mechanical arm. To our knowledge, the published literature contains no quantitative placement of mechanically induced macrophage states on a chemical M1 scale. Two limits of that search should be stated. OpenAlex returned 500 of the 638 and 667 citing works for the two mechanics seeds, so roughly the earliest fifth of those citation lists was not retrieved, and preprint coverage in OpenAlex is incomplete: the claim is made for the published literature rather than for the literature as a whole.

## 2 Materials and Methods

### 2.1 Design

We asked whether mechanically induced macrophage activation engages the same interferon-stimulated gene (ISG) program as chemical M1 polarization. The expected answer was “no.” For that reason, the study was designed as a quantitative equivalence assessment against a within-scale reference, not as a discovery screen: for every mechanical contrast we report the ISG module shift in the same units as the shift produced by LPS + IFN-γ, measured the same way. That way, “no induction” becomes a bounded statement rather than a null result.

The unit of replication is the study, not the sample. All inference below is at the study level. Treating samples as independent would inflate the apparent precision by roughly the average within-study sample size, attributing to replication what is in fact repeated measurement of one experimental system. The cost of the stricter convention: the realised design resolves effects of about 9% of the chemical response, not about 2% (§2.7).

### 2.2 Data sources and selection

Public transcriptomic datasets were retrieved from GEO (NCBI E-utilities ‘esearch’, ‘db=gds’, ‘gse[ETYP]’), ArrayExpress/BioStudies, and ENA/SRA using 48 pre-specified queries (22 GEO, 26 ArrayExpress; 1,971 records). These were supplemented on 2026-11-02 by three further GEO queries, yielding 69 unique series. Query strings and per-query hit counts are given verbatim in Supplementary Table S1; the record-by-record eligibility decisions for all 58 datasets appear in Supplementary Table S3.

After de-duplication, 60 unique accessions were screened on title and sample-level metadata and 58 were verified record by record against five eligibility criteria:

1. Cells: macrophages, monocytes or myeloid cells (primary monocyte-derived macrophages/bone marrow-derived macrophages (MDM/BMDM), microglia, Kupffer cells, tumor-associated macrophages, or the RAW264.7, THP-1 and BV2 lines).
2. Perturbation: an applied or material-imposed mechanical input (substrate stiffness, topography, 3D confinement, stretch, shear, compression, hydrostatic pressure, ultrasound/acoustic oscillation, fluid viscosity, magnetomechanical actuation), or a PIEZO1 agonist as a pharmacological surrogate.
3. Control arm, a within-experiment unstimulated/soft/static control. Designs offering only “mechanics + LPS vs LPS” were excluded at this step. This exclusion removed several otherwise well-powered datasets. The reason is specific to the question: in a design where every arm receives LPS, the ISG module is already at or near its chemically induced ceiling, so a mechanical increment cannot be expressed as a fraction of the chemical response without assuming the very additivity the comparison is meant to test. The same logic excludes designs in which the only control is a different mechanical condition. A 1 kPa versus 50 kPa comparison is informative about the slope of the stiffness response but silent about where either point sits relative to zero.
4. Readout: genome-wide expression (array or RNA-seq; single-cell datasets had to permit myeloid pseudobulk).
5. Availability: a processed matrix, or raw data quantifiable at reasonable cost.

Nine accessions were excluded (SuperSeries container only, n = 4; no expression assay, n = 3; other, n = 2), leaving 49 eligible, of which 33 were quantitatively scored. Twelve studies met criteria 1–5 for the stiffness/applied-force family. GSE274336 and GSE274019 are two series deposited from a single paper; we combined them into one study so that the unit of replication remains the study rather than the deposit. They were scored separately on their own matrices and then combined by inverse-variance weighting (§2.7), not concatenated into a single expression matrix. GSE99445 and GSE99446 are two sub-series of one donor cohort and were likewise treated as two arms of one experiment rather than two studies: this is what allows them to be used as a test of anchor reproducibility (§2.6). Two of these, GSE283034 (pharmacologically softened mammary tumors) and GSE299513 (soft vs stiff mammary tumors), profile whole tumor tissue, so a shift in module score cannot be separated from a shift in myeloid composition. They are pre-specified as a sensitivity set and excluded from the main pool, which comprises K = 10 studies. A further 20 datasets contribute 22 contrasts to the modality projection (§2.8). The 16 unscored datasets lacked an LPS-free arm, were raw-only at disproportionate cost, were access-restricted (GSE291575 returns HTTP 403 for all supplementary files), or could not be reconciled with their deposited record (GSE262146, §4.9). Figure 1 gives the flow diagram.

**Figure 1.**
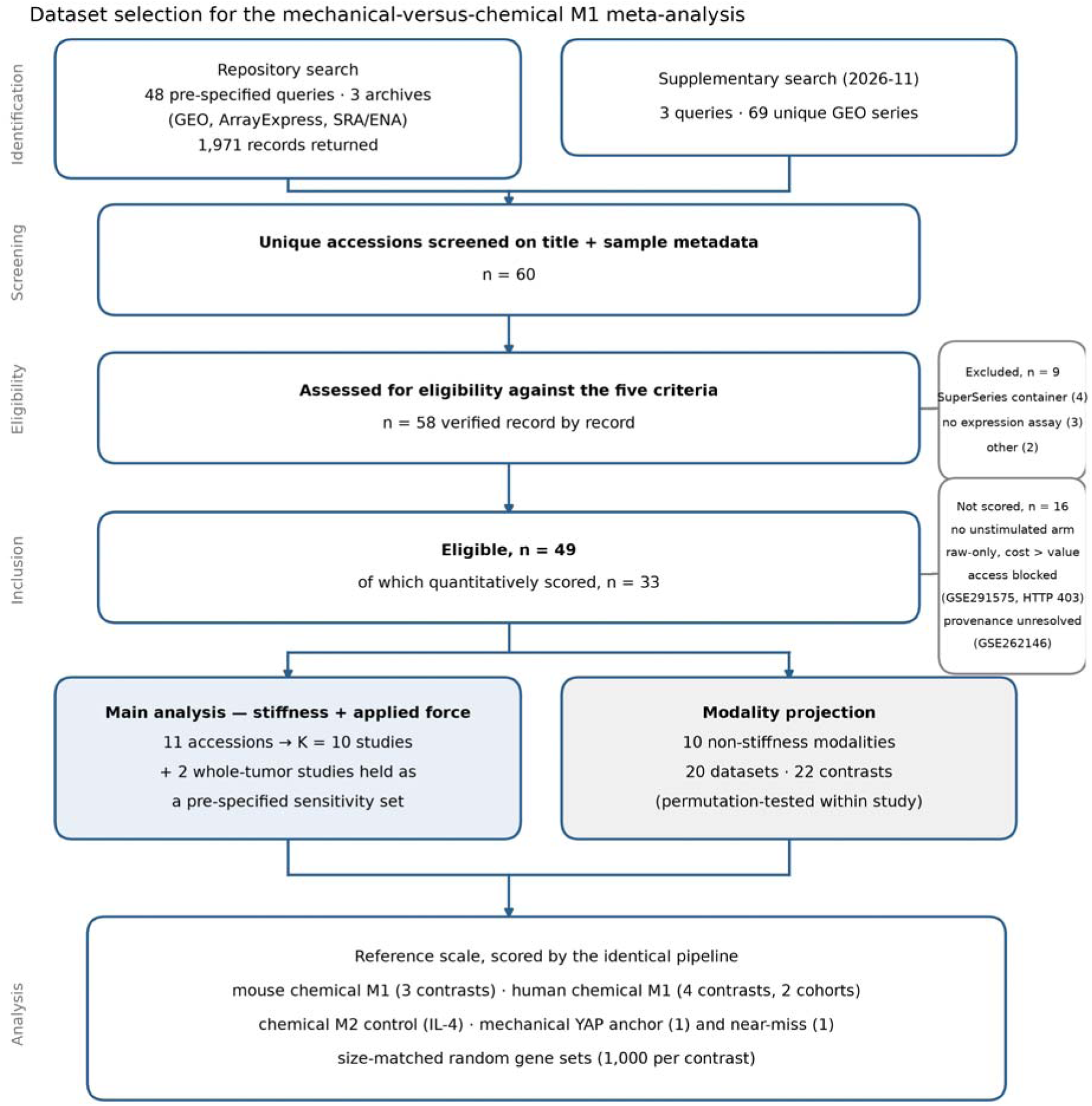
Dataset selection. Flow from the 48 pre-specified repository queries (1,971 records) and three supplementary queries (69 series) through screening, eligibility assessment against the five criteria, and allocation to the main pool (K = 10 studies, with two whole-tumor studies held as a pre-specified sensitivity set) and the modality projection (20 datasets, 22 contrasts). Right-hand boxes give exclusion reasons with counts.

The primary publications of the analysed datasets are cited here so that the provenance of every contrast is traceable. Main-pool studies with a linked publication: GSE206771 (14); GSE239377 (15); GSE273834 (16); GSE36878 (17). Sensitivity-set studies with a linked publication: GSE283034 (18). Modality-projection datasets with a linked publication: GSE125806 (19); GSE133069 (3); GSE143780 (20); GSE186185 (21); GSE190470 (22); GSE236912 (23); GSE273466 (24); GSE273977 (25); GSE274828 (26); GSE275018 (25); GSE290857 (27); GSE303278 (28); GSE304867 (29); GSE314256 (30); GSE344008 (31). GSE262146 (32) appeared in this list in earlier drafts. It was removed from the projection when its provenance could not be re-derived (§4.9); the publication is retained in the reference list because that exclusion is discussed there. The remaining accessions (E-MTAB-7900, E-MTAB-15203, GSE99445, GSE185761, GSE274019, GSE274336, GSE285024, GSE291556, GSE299513, GSE312997, GSE321887, GSE331328) carry no publication linked in the host repository at the time of analysis and are cited by accession only. Their publication status was checked in PubMed by accession and in the repository metadata.

We quantified two datasets from raw reads ourselves, since no processed matrix had been deposited: E-MTAB-7900 (FASTQ from ENA, salmon 2.9.1 (33)) and GSE299513 (author-supplied kallisto transcript abundances aggregated to gene symbols via Ensembl BioMart; 115,095 of 118,489 transcripts mapped). GSE99446 probe sets (custom CDF ‘hugene10st_Hs_ENTREZG’, probe identifiers are Entrez gene IDs) were mapped with NCBI ‘Homo_sapiens.gene_infò. Of 19,700 probes, 19,561 mapped.

### 2.3 Gene modules

The primary module is the ISGF3-specific set of Platanitis et al. (9), taken from their Supplementary Data 1: ISGF3-occupied promoters not occupied by STAT2–IRF9 or STAT1 homodimers. We use this ISGF3-specific set rather than a generic type-I-IFN set because the same work shows IFN-γ also recruits ISGF3 extensively (166 genes in the IFN-γ arm, 359 in the IFN-β arm). A generic “type I ISG” label would make the LPS + IFN-γ reference arm partly self-referential. Human orthologues were mapped with babelgene (34). Two further ISG definitions were carried as sensitivity modules (MSigDB Hallmark IFN-α and Hallmark IFN-γ response (35)), along with the 22-gene YAP/TAZ signature of Wang et al. (36) as a mechanotransduction-pathway module.

Module sizes after mapping (mouse / human): ISGF3-specific 118 / 82; Hallmark IFN-α 92 / 97; Hallmark IFN-γ 193 / 200; YAP/TAZ 22 / 24. A further interferon-stimulated-gene core set (37) was curated but not used in the primary analysis. All 13 curated sets, with their source DOI and supplementary-table location, are provided as Supplementary Table S2.

Effector-arm modules. The interferon arm is only one part of the chemical M1 program. Three NF-κB/AP-1 effector modules were added, so the same contrasts can be read on both arms through one pipeline: MSigDB Hallmark TNFA_SIGNALING_VIA_NFKB (35) (200 / 200 genes, mouse / human), Hallmark INFLAMMATORY_RESPONSE (200 / 201), and a seven-gene field panel assembled from the markers most often used to assert “mechanically induced M1” (mouse *Nos2, Tnf, Il6, Il12b, Il1b, Ccl2, Cxcl1*; human *TNF, IL6, IL12B, IL1B, CCL2, CXCL8*, with *NOS2* omitted because human macrophages express it poorly; 7 / 6 genes). Mouse sets were taken from msigdbr and orthologue-mapped with babelgene (34) by the same route as the interferon modules. The Hallmark modules retrieved in this step reproduce the sets used in §2.3 exactly (97 / 97 and 200 / 200 genes), confirming the stored interferon sets.

The two arms are nearly disjoint at the gene level: the ISGF3-specific set and Hallmark TNFA_SIGNALING_VIA_NFKB share 5 of 82 human genes. A dissociation between them is a statement about two largely non-overlapping gene populations, not an arithmetic consequence of shared membership.

One asymmetry in construction must be stated plainly: it is not removable with currently distributed resources. The interferon module is defined by measured ISGF3 promoter occupancy (9). No equivalent RELA occupancy set exists in the GTRD collection distributed with msigdbr; the only related entry is NFKBIA target genes (1,348 genes), which is both the wrong transcription factor and far too large to behave as a module. The effector arm is defined by expression-derived Hallmark sets, the interferon arm by occupancy. We did not attempt to equalise this, and the two modules are not equally specific.

### 2.4 Expression processing and module scoring

Counts were converted to log CPM, TPM/FPKM and normalized array intensities to log (x + 1), and already-logged matrices were used as supplied. Identifiers were mapped to gene symbols; duplicate symbols were collapsed to the row of highest mean expression.

One ranking background is used for every module score in this work. Because singscore ranks genes within each sample, the size of the retained gene universe sets the scale of every score difference: a module shift computed against a smaller background is proportionally larger, so mixing backgrounds within one paper would make percentages mutually incomparable. That single background is the full symbol-mapped gene universe with no detection filter, for example 33,043 genes for GSE274336 and 25,850 for the GSE274020 anchor. It applies to the main pool (§3.2–§3.4), the chemical and mechanical anchors (§2.6), the modality projection (§3.5), the magnetomechanical contrasts (§3.7) and both arms of §3.9. A common default is to drop genes detected in fewer than max (2, 0.25 × n_samples) samples; its effect here is not small. Doing so inflates the shift of an RNA-seq contrast by roughly 1.65-fold. Array data, which has no zero-valued probes, behaves differently: the same filter is close to a no-op, which is why array-based quantities come out numerically identical either way while sequencing-based ones do not.

Differential-expression counts are computed on the detection-filtered matrix instead, for a reason specific to that statistic. A rank-based score is well defined when many genes are tied at zero, but an empirical-Bayes variance model is not. On the unfiltered matrices used here, 18–36% of genes have exactly zero variance across samples (18.3% for the GSE275018 myeloid pseudobulk, 22.6% for GSE274336, 35.6% for GSE304867), and limma issues an explicit warning that its moderated variances are unreliable in that regime. Every differential-expression count in this paper is taken from the matrix after the detection filter, where no gene has zero variance. The two statistics are preprocessed according to their own requirements, not forced onto a common matrix: per-contrast background sizes for both are reported in Supplementary Tables S10 and S15.

Per-sample module activity was computed with singscore (v1.26.0) ‘simpleScorè (38), using the up-set form with ‘centerScore = TRUÈ: the mean rank of the module genes normalized to [0, 1] against the attainable range and centred at 0.5. This is a rank-based, single-sample statistic. It requires no cross-study normalization and no batch correction, which is why none was applied anywhere in this work.

### 2.5 Contrast-level effect, within-study permutation, and random-set calibration

For each contrast the effect is the difference in module score, Δ = mean(mechanical) − mean(control), computed within donor or batch stratum and then averaged across strata. For donor-paired designs, the per-donor difference is averaged, and its standard error is the SD of those differences divided by √n_donors.

Two calibrations accompany every Δ:

- **Within-study label permutation.** All label assignments respecting the stratification were enumerated exhaustively when the permutation space was ≤ 20,000, otherwise 20,000 were sampled. The two-sided p is the fraction of permuted |Δ| at least as large as observed. The attainable minimum is set by the design: a donor-paired 6-donor contrast cannot go below 2/2 = 0.031, a 2-vs-2 contrast cannot go below 1/3. We report attainable minima alongside p values throughout.
- **Size-matched random gene sets**. 1,000 random gene sets of the same size were drawn from the detected background and scored identically, giving z = (Δ − mean_random)/SD_random and an empirical p. This separates “the module moved” from “everything moved.”

The two calibrations answer different questions, and neither substitutes for the other. The permutation test asks whether the observed separation between the mechanical and control arms exceeds what the same samples produce under relabelling; it is sensitive to sample size in a way that is explicit rather than modelled. The random-set test asks something else: whether the module moved more than an arbitrary gene set of the same size drawn from the same expression background. This separates a module-specific response from a global shift in rank structure, the failure mode that most often produces a spurious pathway result in small transcriptomic contrasts. One compression contrast passes the permutation test but fails the random-set test (§3.6). It has moved the module no more than chance gene sets of equal size, and we do not treat it as evidence of a module-specific effect.

Differential expression for each contrast was computed with limma (v3.62.1) (39) via ‘lmFit’/‘eBayes (trend = TRUE)’ on the same design, and is reported as the number of genes at FDR < 0.05. This is an amplitude check: it establishes that a contrast with an unmoved ISG module is nevertheless a transcriptionally real perturbation.

### 2.6 Reference anchors

Chemical M1 anchors were scored by the identical pipeline. In mouse: GSE69607 M1 (LPS + IFN-γ) vs M0, Δ_ISGF3 = +0.0392 (g = 3.93); GSE253423 LPS 4 h, +0.1380 (g = 10.53); GSE274020 LPS 6 h, +0.1245 (g = 2.65). The mouse anchor used throughout is the median, 0.1245. In human, GSE46903 gives M1 vs M0 +0.0770 and IFN-γ 72 h +0.0760. Both were re-derived here from the deposited series matrix (19,262 symbol-mapped probes): the values carried in earlier drafts could not be traced to a surviving computation. GSE99445 within-study IFN-γ (8 paired donors) gives +0.1002, the human anchor.

Anchor reproducibility was itself tested, since the whole analysis is expressed as a fraction of it. Two sub-series of the same cohort (GSE99445, porous-Ti arm, 8 donors; GSE99446, polished-Ti arm, 6 donors) give ISGF3 +0.10017 vs +0.10166: agreement to within 1.5%. The same comparison for the YAP module gives +0.02684 vs +0.01099, a 2.44-fold discrepancy. We treat the ISGF3 ruler as reliable and attach an explicit factor-2.4 uncertainty to every YAP-module quantity.

Mechanical anchors for the YAP module were identified by screening all YAP contrasts for within-study permutation p < 0.05 and random-set p < 0.05. One contrast qualifies: GSE99445, porous Ti vs plastic (8 paired donors, Δ = +0.01085, p_perm = 0.039, p_rand = 0.035, 40 DE genes). The nearest miss is E-MTAB-15203, Yoda1 vs DMSO (human MDM, 6 paired donors, Δ = +0.01707, 4,421 DE genes): its permutation p of 0.063 is the second attainable level for a six-donor paired design and its random-set p is 0.108. We carry it as a weaker second candidate and report margins against both. A compression contrast (GSE304867) was rejected on both tests (p_perm = 0.303, p_rand = 0.641).

Effector-arm anchors require LPS, and an early time point. The IFN-γ anchors above cannot serve the effector modules. Scored on the same two sub-series that validate the interferon ruler to within 1.5%, the effector modules do not replicate: TNFA/NF-κB gives +0.02917 (GSE99445) vs +0.01256 (GSE99446), a 2.32-fold discrepancy, and the field panel gives +0.02183 versus −0.05103, a sign flip (Figure 2B). This makes sense: IFN-γ signals through STAT1 and does not itself engage NF-κB, so an IFN-γ contrast offer no stable effector signal to anchor against.

**Figure 2.**
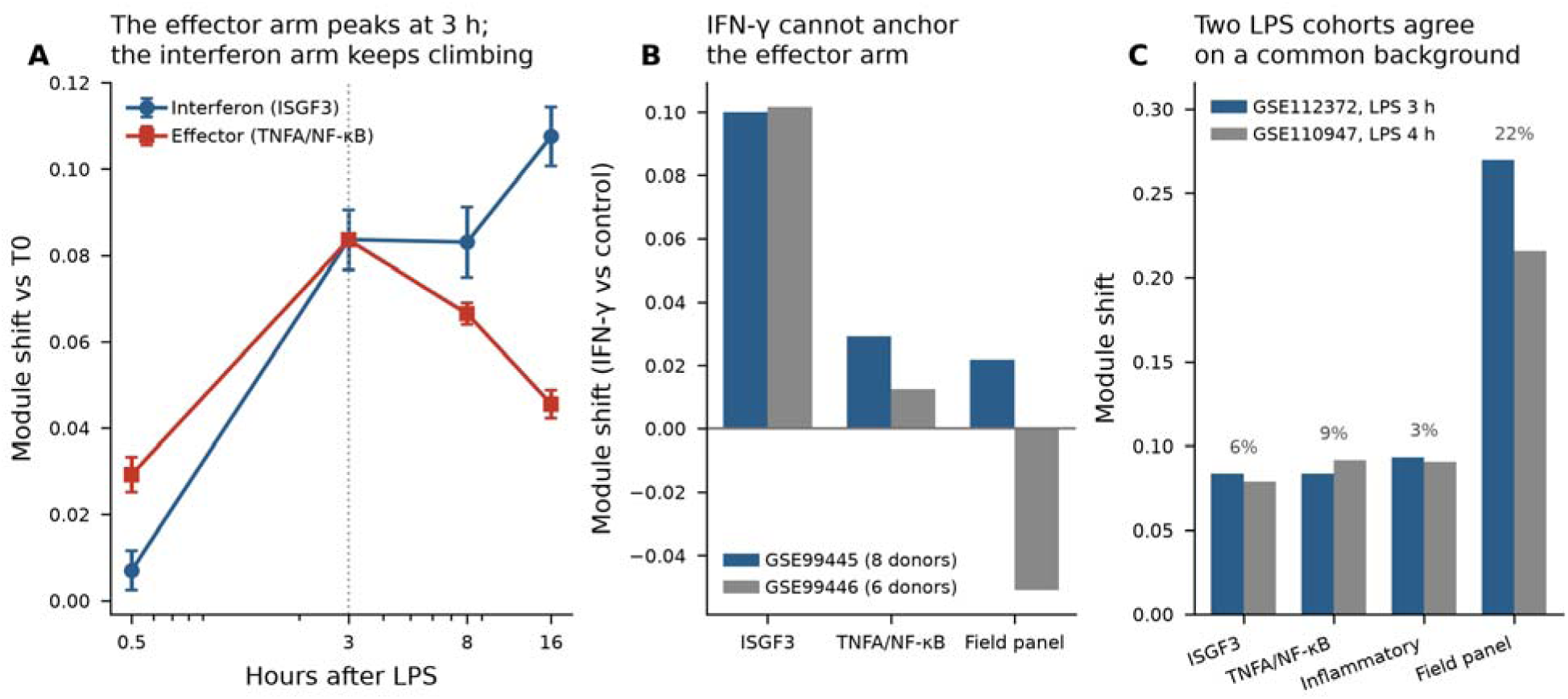
Building a reference scale for the effector arm. (A) Time course of human monocyte-derived macrophages stimulated with LPS (GSE112372, four fully paired donors). The NF-κB effector modul peaks at 3 h and decays by 16 h; the ISGF3-specific interferon module rises monotonically instead. At 0.5 h the effector arm has already moved and the interferon arm has not. Error bars are standard errors of the per-donor differences. (B) The IFN-γ contrast cannot anchor the effector arm. Two sub-series of one donor cohort agree to 1% on the interferon module but differ 2.32-fold on TNFA/NF-κB, and flip sign on the seven-gene field panel. (C) With an early LPS contrast the ruler holds. On a common background, the sensitivity cohort (GSE110947, LPS 4 h) and the primary cohort (GSE112372, LPS 3 h) agree to 6% on the interferon module, 9% on TNFA/NF-κB, and 3% on the inflammatory module; only the six-gene field panel differs materially (22%).

Two human LPS cohorts were added. In GSE112372 (40) (four fully paired donors; T0 control plus LPS at 0.5, 3, 8 and 16 h) the two arms resolve in time (Figure 2A). At 0.5 h the effector arm has already moved (TNFA/NF-κB +0.0292), while the interferon arm has not (+0.0071). The effector arm peaks at 3 h (+0.0836, SE 0.0009) and decays to +0.0456 by 16 h; the interferon arm keeps climbing instead, plateauing between 3 and 8 h (+0.0837 and +0.0831) before reaching +0.1076 at 16 h and overtaking the effector arm by a factor of 2.4. The 3 h contrast is the primary human effector anchor (ISGF3 +0.0837, TNFA/NF-κB +0.0836, inflammatory +0.0935, field panel +0.2697): the effector arm is at its maximum here, and the two arms are still level, so neither is scaled against a reference already past its own peak. GSE110947 (five paired donors, LPS 4 h) is the sensitivity anchor. On a common background the two cohorts agree directly (Figure 2C): interferon +0.0790 vs +0.0837 (6% apart), TNFA/NF-κB +0.0916 vs +0.0836 (9%), inflammatory +0.0908 vs +0.0935 (3%). Only the six-gene field panel differs materially (+0.2154 vs +0.2697, 22%), which is what one expects of a module that small. The effector-to-interferon ratio agrees between the cohorts to 15% (TNFA/NF-κB), 3% (inflammatory) and 18% (field panel).

Two consequences are carried through the analysis. First, the human effector percentages carry a between-cohort uncertainty of order 10% for the two large modules and 22% for the six-gene panel. That is small enough to report absolute percentages directly, but we also give the within-contrast ratio between the arms wherever the distinction could change a reading. Second, an effector anchor must be taken from an early LPS window. The widely used 72 h arms of GSE46903 lie well past it, and its two LPS preparations disagree there even in sign (TNFA/NF-κB +0.0485 for standard LPS vs −0.0296 for ultrapure LPS).

Anchor sets used for the two arms. All modules are scored on one background (§2.4), so both arms can be divided by anchors measured in a single pass: interferon percentages mean the same thing everywhere in the paper. The mouse interferon anchor is 0.1245 in §3.2–§3.8 and in §3.9 alike. The effector anchors come from the LPS contrasts of this section, not the IFN-γ contrasts: IFN-γ signals through STAT1 and does not itself engage NF-κB. In mouse (GSE274020, LPS 6 h), TNFA/NF-κB is 0.0976, inflammatory is 0.1047 and field panel is 0.3539; in human (GSE112372, LPS 3 h), interferon is 0.0837, TNFA/NF-κB is 0.0836, inflammatory is 0.0935 and field panel is 0.2697. The human interferon arm of §3.9 is scaled against LPS rather than against the IFN-γ anchor of §2.6. This is the appropriate comparator when the two arms are set side by side, and it is the only place where a §3.9 interferon percentage is referred to a different contrast from the one used in §3.2–§3.8. All anchors are given in Supplementary Table S9.

### 2.7 Meta-analysis

Where one study contributed two deposited series (GSE274336 and GSE274019, study S1), each series was scored on its own matrix and the two estimates were combined by fixed-effect inverse-variance weighting before entering the pool. The study enters with a single Δ and a single SE: Δ_S1 = +0.004305, SE 0.003947, from Δ = −0.002544 (SE 0.005099; 12 vs 12, two harvest-time strata) and Δ = +0.014545 (SE 0.006235; 3 vs 3, wild-type arm only). Study-level Δ and SE were then pooled by DerSimonian–Laird random effects (41); heterogeneity is reported as τ² and I². Inverse-variance weights are extremely unequal here, so we also report the Kish effective number of studies (42), n_eff = (Σw)²/Σw², which for the main pool (K = 10) is 5.72 under random-effects weights: the honest denominator for any statement about replication breadth.

The Kish correction matters more here than in a typical meta-analysis. Sample sizes in this literature range from two per arm to twelve per arm, so inverse-variance weights span roughly two orders of magnitude, and a nominal K of ten can be dominated by two or three well-powered studies. Reporting K alone would overstate the breadth of replication. Reporting n_eff alongside it states how many studies the estimate effectively rests on.

The primary test is a study-level sign-flip permutation over all 2^K = 1,024 sign configurations, whose attainable two-sided minimum is 2/2¹ = 1.95 × 10 ³. We prefer this to a parametric test on the pooled estimate: it makes no distributional assumption about the between-study effects, and its resolution limit is transparent. With ten studies no configuration can produce a two-sided p below 1.95 × 10 ³. With three studies, the human stratum, none can produce one below 0.25. Quoting a permutation p without its attainable minimum invites a reader to interpret p = 0.25 as weak evidence when it is in fact the strongest result the design can return.

Equivalence testing and the origin of the margin. Two one-sided tests (TOST) were applied to the random-effects estimate with df = K − 1, against margins expressed as a percentage of the species-matched chemical anchor. The ±10% margin is post hoc: it was not pre-registered and does not represent a clinical or biological significance threshold. It was chosen because the realised design supports it. The minimum detectable effect at 80% power is 8.8% of the anchor (§ below). The next tighter margin is not supported at all: in the main pool ±5% is not met (p = 0.058) and fails in 7 of 10 leave-one-out refits, while ±10% holds in all 10 (max p = 0.0139). The estimate (μ and its 95% CI as a percentage of the anchor) is the primary quantity reported in the abstract, with TOST results given in support.

Power and minimum detectable effect were obtained by Monte Carlo over the realised SE structure and τ²; we simulated y ∼ N(δ, SE² + τ²) and applied the same sign-flip test to each replicate. MDE (80%) was located by bisection.

Robustness. We reran every reported quantity using leave-one-study-out refits, three independent ISG module definitions, and species-stratified pooling.

Small-study effects. Egger’s regression (weighted least squares of Δ on SE, weights 1/SE²) and Begg’s rank-correlation test were applied to each module separately. Both have low power at K = 10, so we additionally report the raw correlation between Δ and SE.

### 2.8 Modality projection

Datasets satisfying criteria 1–5 but imposing a mechanical input other than substrate stiffness were scored identically. These are reported as a projection across 10 modalities (20 datasets, 22 contrasts), not pooled into the main estimate. Where pooling within a modality family was performed, it is reported with its I². Pooled values for families with I² > 85% are not interpreted as modality effects.

### 2.9 Validation strategy

Because this is a secondary analysis of archived transcriptomes, no new primary measurement can validate the estimate. We pre-specified five internal validations: each addresses a distinct way in which a null result of this kind can be wrong. We report all five regardless of outcome.

1. Scale validation. The chemical reference is scored by the identical pipeline and must move the module before any mechanical absence is interpreted (§3.1). We test its reproducibility by measuring the same stimulus in two independent sub-series of one donor cohort.
2. Gene-set validation. The primary ISGF3 module is replaced by two independently derived definitions, Hallmark IFN-α and Hallmark IFN-γ, and the conclusion must survive all three.
3. Specificity validation. Every contrast is scored against 1,000 size-matched random gene sets drawn from the same detected background, distinguishing “the module did not move” from “nothing moved”.
4. Amplitude validation. Each contrast reports its differential-expression count, establishing that a contrast with an unmoved module is nevertheless a transcriptionally real perturbation.
5. Replication validation. Leave-one-study-out refitting, species stratification and sequential expansion of the pool (K = 8 → 10 → 12) test whether the estimate depends on any single study or species. The Monte-Carlo power analysis states the effect size the design can resolve.
6. Two-arm validation. Every contrast is also scored on NF-κB effector modules through the identical pipeline, which distinguishes an unmoved interferon module from an unmoved cell. The effector ruler is itself validated before use. We report the anchor contrast on which it fails (IFN-γ) as well as the one on which it holds (early LPS) (§2.6, Figure 2).

For an equivalence claim these are the appropriate validations: a single additional wet-laboratory experiment cannot falsify “no net induction across studies,” while repetition across independent studies can. That is what the sequential expansion of the pool does. The validations fail in two places: the sign of the pooled estimate is not robust to leave-one-out, and equivalence at a ±5% margin is not supported (§3.4).

### 2.10 Software

R 4.4.3 (singscore 1.26.0, limma 3.62.1, GSEABase 1.68.0); Python 3.11.18 (numpy 2.4.6, scipy 1.17.1, pandas 2.3.3, statsmodels 0.15.0, matplotlib 3.11.2); salmon 2.9.1; scanpy (43) for single-cell pseudobulk (Leiden clustering (44), resolution 1.0). Analysis code and all intermediate tables are provided as Supplementary Data and in the repository named in the Data Availability Statement.

## 3 Results

### 3.1 The ruler works before the measurement is attempted

Any statement of the form “mechanics does not induce ISGs” is only as good as the scale it is denied on. We first confirmed the ISGF3 module is detectable, large, and reproducible when it is genuinely engaged.

Chemical M1 moved the module strongly and consistently across platforms and cell sources. In mouse: GSE69607 M1 (LPS + IFN-γ) vs M0, Δ_ISGF3 = +0.0392 (Hedges’ g = 3.93); GSE253423 LPS 4 h, +0.1380 (g = 10.53); GSE274020 LPS 6 h, +0.1245 (g = 2.65) (Figure 3A). In human: GSE46903 M1 vs M0 +0.0770 (g = 4.32) (§2.6) and IFN-γ 72 h, 7 paired donors, +0.0760 (g = 3.07); GSE99445 within-study IFN-γ, 8 paired donors, +0.1002 (Figure 3A). The module is measurable on microarrays and RNA-seq, in cell lines and in primary BMDM and MDM. A failure to detect it on the mechanical side is not a pipeline failure.

**Figure 3.**
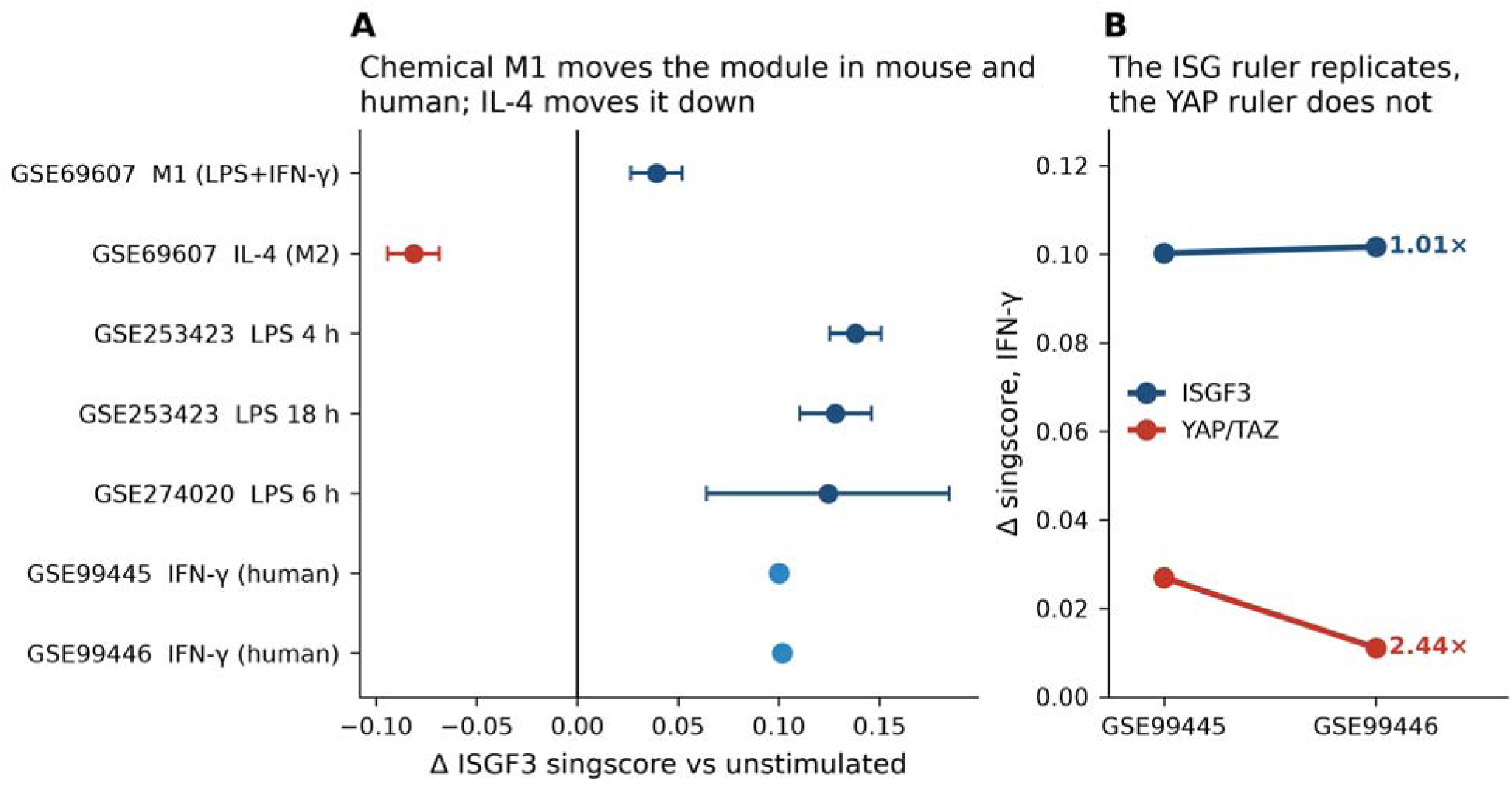
We establish the reference scale before interpreting any mechanical absence. (A) ISGF3 module shift for the mouse chemical anchor contrasts (dark blue) and the two human IFN-γ anchor sub-series (light blue); bars are 95% confidence intervals where a within-study standard error is available. The two human points carry none, since only the donor-paired point estimate was retained. IL-4 (M2) polarization (red) moves the same module in the opposite direction. The remaining human anchors (GSE46903) are given in §2.6. (B) Anchor reproducibility: two sub-series of one donor cohort, differing only in implant surface, give IFN-γ anchors agreeing to within 1.5% on the ISGF3 module but differing 2.44-fold on the Wang 22-gene YAP/TAZ module. The ISG ruler is reliable at the precision used here. The YAP ruler is not.

The ruler is also reproducible. Two sub-series of one donor cohort, differing only in implant surface, give an IFN-γ anchor of +0.10017 (GSE99445, 8 donors) and +0.10166 (GSE99446, 6 donors): agreement to within 1.5% (Figure 3B). The same comparison for the YAP/TAZ module gives +0.02684 vs +0.01099, a 2.44-fold discrepancy (Figure 3B) (§2.6). The ISG scale is trustworthy at the precision we use it. The YAP scale is not, and every YAP quantity below carries that factor explicitly.

Specificity runs in the expected direction: IL-4 (M2) polarization moves the same module down, by −0.0813 in GSE69607 (Figure 3A).

### 3.2 Across ten independent studies the mechanical shift is 0.9% of chemical M1

Pooling the ten studies of the stiffness/applied-force family by random effects gives μ = −0.00111 singscore units, 95% CI [−0.00776, +0.00555] = −0.89% of the chemical M1 anchor, 95% CI −6.2% to +4.5%, with τ² = 2.73 × 10□□and I² = 41.1% (Figure 4). The study-level sign-flip permutation is p = 0.711 against an attainable minimum of 2/2¹□= 1.95 × 10□³: the pooled estimate is indistinguishable from zero in a test that had the resolution to say otherwise. Equivalence is established at a ±10% margin (TOST p = 0.0019) and at ±20% (p = 1 × 10□□), but not at ±5% (p = 0.058).

**Figure 4.**
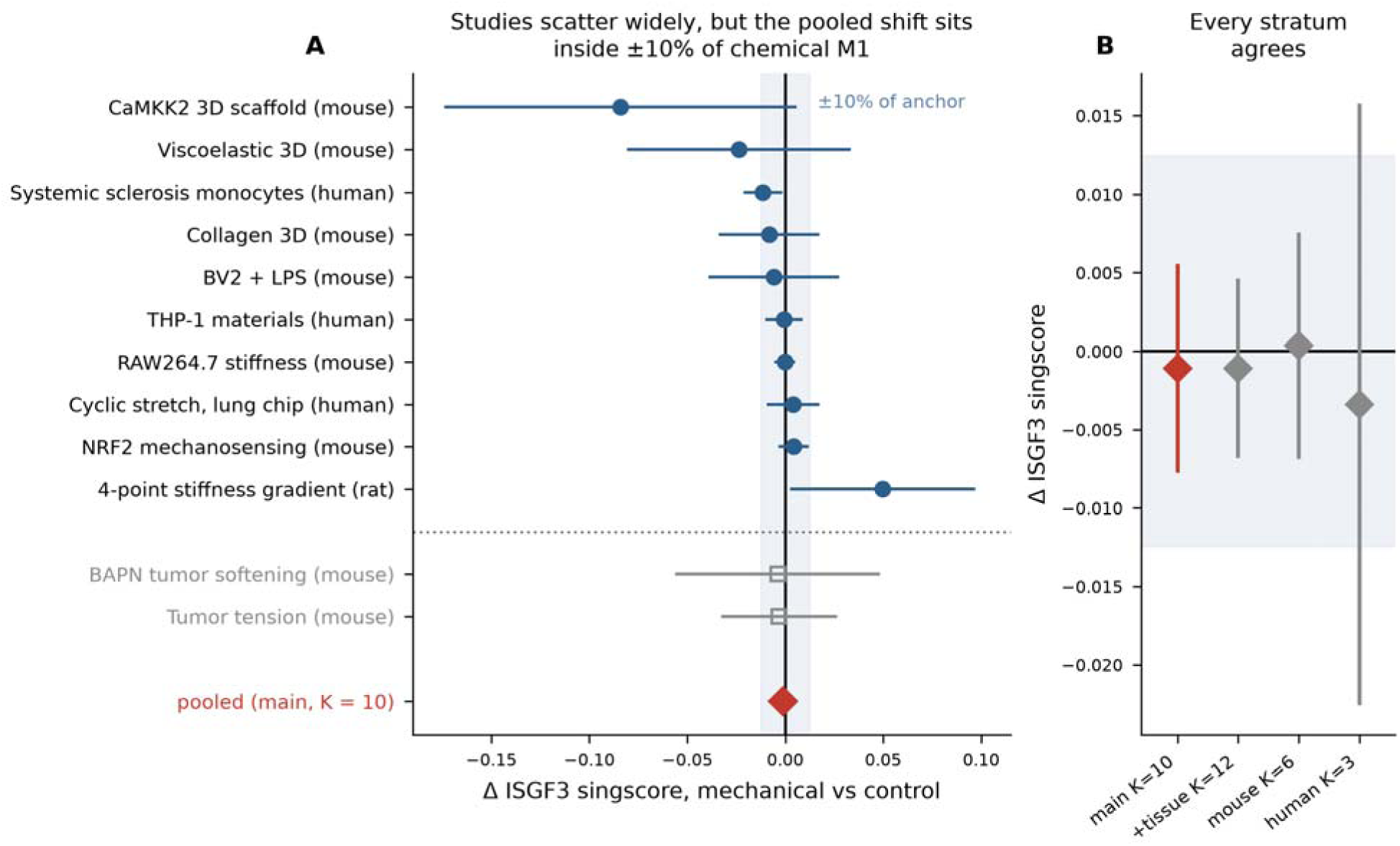
Pooled mechanical shift. (A) Study-level ISGF3 shifts with 95% confidence intervals for the ten main-pool studies (filled circles) and the two sensitivity-set whole-tumor studies (open squares, below th dotted rule). The shaded band is ±10% of the species-matched chemical M1 anchor; the red diamond marks the random-effects pooled estimate. (B) The same estimate recomputed in each stratum and with the sensitivity set added: the answer does not depend on the partition.

The ten study-level estimates are not uniformly small. They are scattered and individually imprecise, which is the pattern a genuine null produces in a literature of small experiments (Figure 4A). Ordered by effect, they run from −0.0841 ± 0.0458 (CaMKK2-dependent stiffness response in 3D scaffolds, GSE273834) and −0.0237 ± 0.0291 (matrix viscoelasticity, GSE206771) at the negative extreme, through −0.0114 ± 0.0051 (systemic-sclerosis monocytes, GSE321887), −0.0083 ± 0.0131 (3D collagen, 7 vs 2 mg mL ¹, GSE239377), −0.0059 ± 0.0170 (BV2 microglia, GSE291556), −0.0007 ± 0.0049 (THP-1 on engineered materials, GSE301422) and −0.0002 ± 0.0027 (RAW264.7 stiffness, GSE36878) near zero, to +0.0039 ± 0.0068 (human cyclic stretch, GSE312997), +0.0043 ± 0.0039 (NRF2 mechanosensing, GSE274336/GSE274019) and +0.0498 ± 0.0241 (four-point stiffness gradient, E-MTAB-7900) at th positive extreme (Figure 4A). Only two of the ten have confidence intervals excluding zero, and they point in opposite directions: the systemic-sclerosis monocytes negative, the rat stiffness gradient positive. Neither survives as a pooled signal. The spread is consistent with τ² = 2.73 × 10□□, a between-study standard deviation of 0.0052 singscore units, or 4.2% of the anchor.

The effective number of studies is smaller than ten: Kish n_eff = 5.72 under random-effects weights. All statements about replication breadth below use that denominator. Two of the ten studies, the RAW264.7 stiffness series and the THP-1 materials series, together carry a majority of the fixed-effect weight. The random-effects weighting used throughout flattens this, which is why the random-effects n_eff is quoted rather than the fixed-effect value.

The strongest single piece of evidence is a contrast that worked. GSE274336 is the most transcriptionally responsive mechanical comparison in the set: 4,318 genes at FDR < 0.05 (n = 12 vs 12, stratified b harvest time) (Figure 5). Its ISGF3 shift is −0.0025, sitting in the middle of the distribution of 1,000 size-matched random gene sets (z = −0.98, p_random = 0.52), or −2.0% of chemical M1. A perturbation that demonstrably rewires four thousand genes leaves this module untouched. The full five-module result for this contrast, with its detected gene background of 33,043 genes, is Supplementary Table S15. The same pattern recurs in the one human cyclic-stretch dataset. GSE312997 moves 1,706 genes and shifts ISGF3 by +0.0039, +3.9% of the human anchor, z = +0.83.

**Figure 5.**
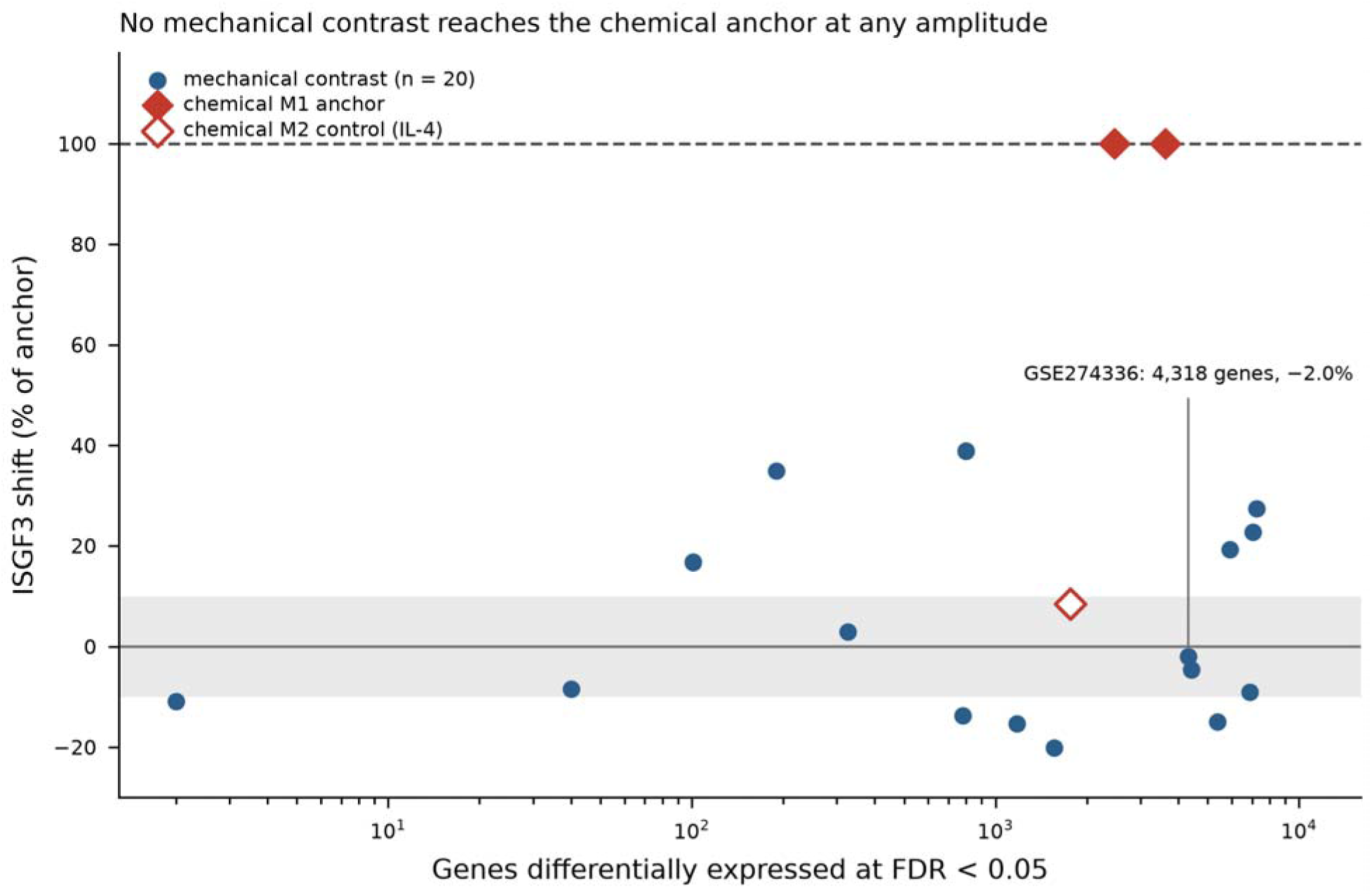
Transcriptional amplitude does not buy interferon engagement. Each point is one contrast, plotted as ISGF3 shift (percentage of the species-matched chemical anchor) against the number of genes differentially expressed at FDR < 0.05. Mechanical contrasts (blue, n = 20) span −21.0% to +38.8%, and 12 of the 20 fall outside the ±10% band, with absolute shift rising weakly with amplitude (Spearman ρ = 0.38, p = 0.097). None, however, approaches the chemical anchors (red diamonds, 100% by construction) at any amplitude. The open red diamond is the chemical M2 control (IL-4). Contrasts with zero genes differentially expressed at FDR < 0.05 cannot be placed on the logarithmic amplitude axis and are omitted; their effect sizes appear in Supplementary Tables S4 and S10.

### 3.3 The answer does not depend on which ISG definition is used

Three independently derived definitions, two curated Hallmark sets and one built from ChIP-defined ISGF3 occupancy, agree on both sign and magnitude (Table 1). The conclusion is a property of the data, not of the gene list.

**Table 1.**
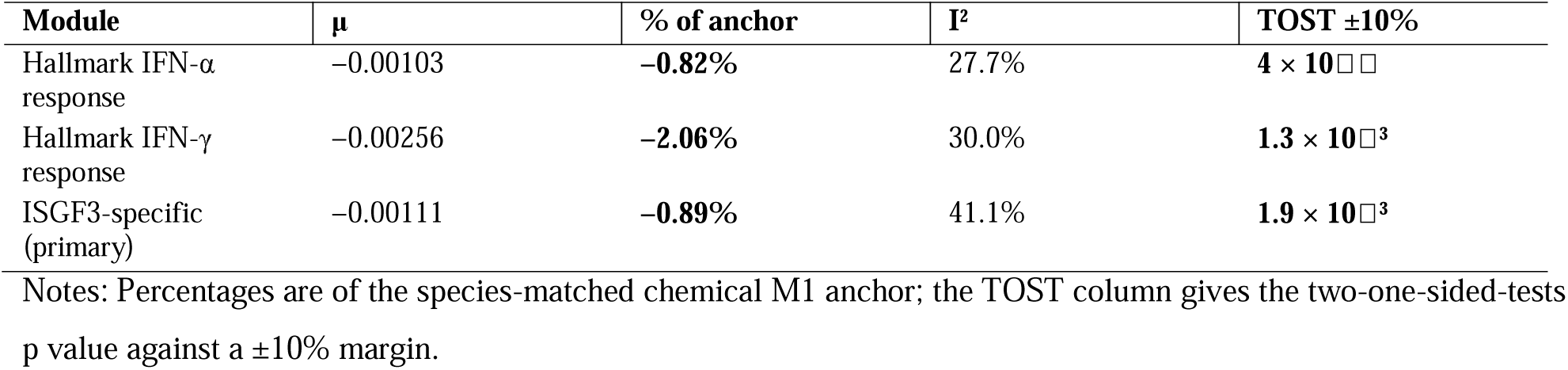
Pooled mechanical shift under three independently derived interferon-stimulated gene definitions (K = 10 main pool, random effects).

### 3.4 Robustness

Leave-one-out. Equivalence at ±10% survives all ten refits (worst p = 0.0139) (Figure 6A). The point estimate ranges from −2.01% to +0.79% of the anchor. The sign is not robust: removing the systemic-sclerosis monocyte study (GSE321887) flips the pooled estimate positive (+0.79%). We make no claim that mechanics suppresses ISGs. It does not induce them at a detectable fraction of the chemical response.

**Figure 6.**
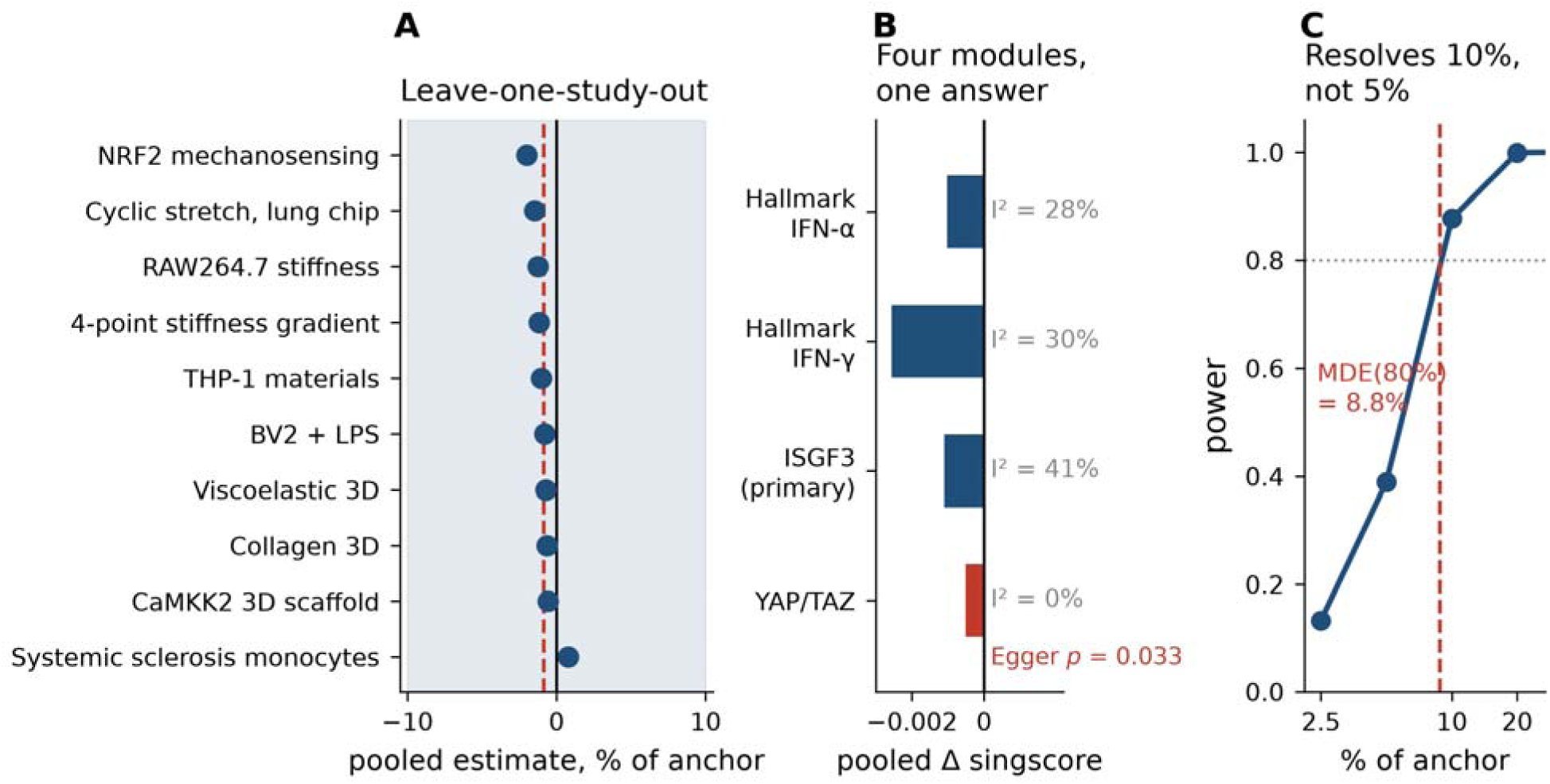
Robustness of the pooled estimate. (A) Leave-one-study-out refits: the dashed red line is the full-pool estimate, and the shaded band is ±10% of the anchor. The magnitude is stable, but the sign is not: removing the systemic-sclerosis monocyte study flips it positive. (B) Pooled estimate for all four modules with heterogeneity and Egger’s test; only YAP/TAZ shows a detectable small-study effect. Study names in (A) are abbreviated; the species of each is in Supplementary Table S4. (C) Monte-Carlo power of the study-level sign-flip test against the realised standard-error structure and τ². The minimum detectable effect at 80% power is 8.8% of the anchor, which is what sets the declared ±10% margin.

Whole-tissue sensitivity set. Two in-vivo whole-tumor studies were excluded from the main pool: composition and macrophage-intrinsic response cannot be separated in bulk tissue (GSE283034, Δ = −0.0039 ± 0.0267; GSE299513, Δ = −0.0032 ± 0.0151). Adding them back changes the pooled estimate from −0.89% to −0.89%, identical to two decimal places, and tightens the CI to [−5.5%, +3.7%] (Figure 4).

Species. In mouse (K = 6) the pooled shift is +0.28% of the mouse anchor (95% CI −5.5% to +6.1%) with I² = 12.0%; equivalence holds at ±10% (p = 0.0038) (Figure 4B). In human (K = 3) it is −3.38% of the human anchor (I² = 48.8%); equivalence holds at ±20% (p = 0.032) but not at ±10% (p = 0.137) (Figure 4B). The human stratum is the binding limitation of this work. A study-level permutation over three studies cannot return a p below 0.25, so the human statement rests on the parametric TOST with df = 2: read it as directional.

Small-study effects. Egger’s regression and Begg’s test are null for all three ISG modules (Egger p = 0.43–0.58; Begg p = 0.29–0.60) (Figure 6B). This should not be over-read: at K = 10 these tests have little power, and the raw correlation between Δ and SE is −0.58 for ISGF3. The least precise studies lie systematically on the negative side. This is the same phenomenon as the non-robust sign in the leave-one-out analysis; read the two together rather than as independent reassurances.

Power. Against the realised SE structure and heterogeneity, the design has 87.7% power to detect a true effect of 10% of the chemical anchor and 38.9% at 5%; the minimum detectable effect at 80% power is 0.0109 singscore units = 8.8% of the anchor (Figure 6C). The ±10% margin matches what the data can resolve. The failure at ±5% is a statement about the evidence base, not about the biology.

### 3.5 Mechanical modalities are not a flat line, but none reaches the chemical scale

Projecting the same score across ten non-stiffness modalities (20 datasets, 22 contrasts) shows that mechanics is not inert with respect to ISGs (Figure 7). The shifts span −0.0251 to +0.0696 and separate into two groups. Lowering the module: in-vivo fluid shear during cardiopulmonary bypass (−21.0% of anchor), PEG hydrogel implantation (−20.2%), a rotating magnetic field applied to lysosome-loaded macrophages (−15.4%), ultrasound on a piezoelectric membrane (−15.0%, within-study p = 0.029), ultrasound alone (−10.9%), compression (−9.0%, p = 0.004). Raising it: in-vivo cyclic stretch of skin (+55.9%), mechanical ventilation (+38.8%), 5 kHz acoustic oscillation (+34.9%), cyclic hydrostatic pressure (+31.1%), fluid viscosity (+27.4%), ultrasound of human macrophages (+22.7%).

**Figure 7.**
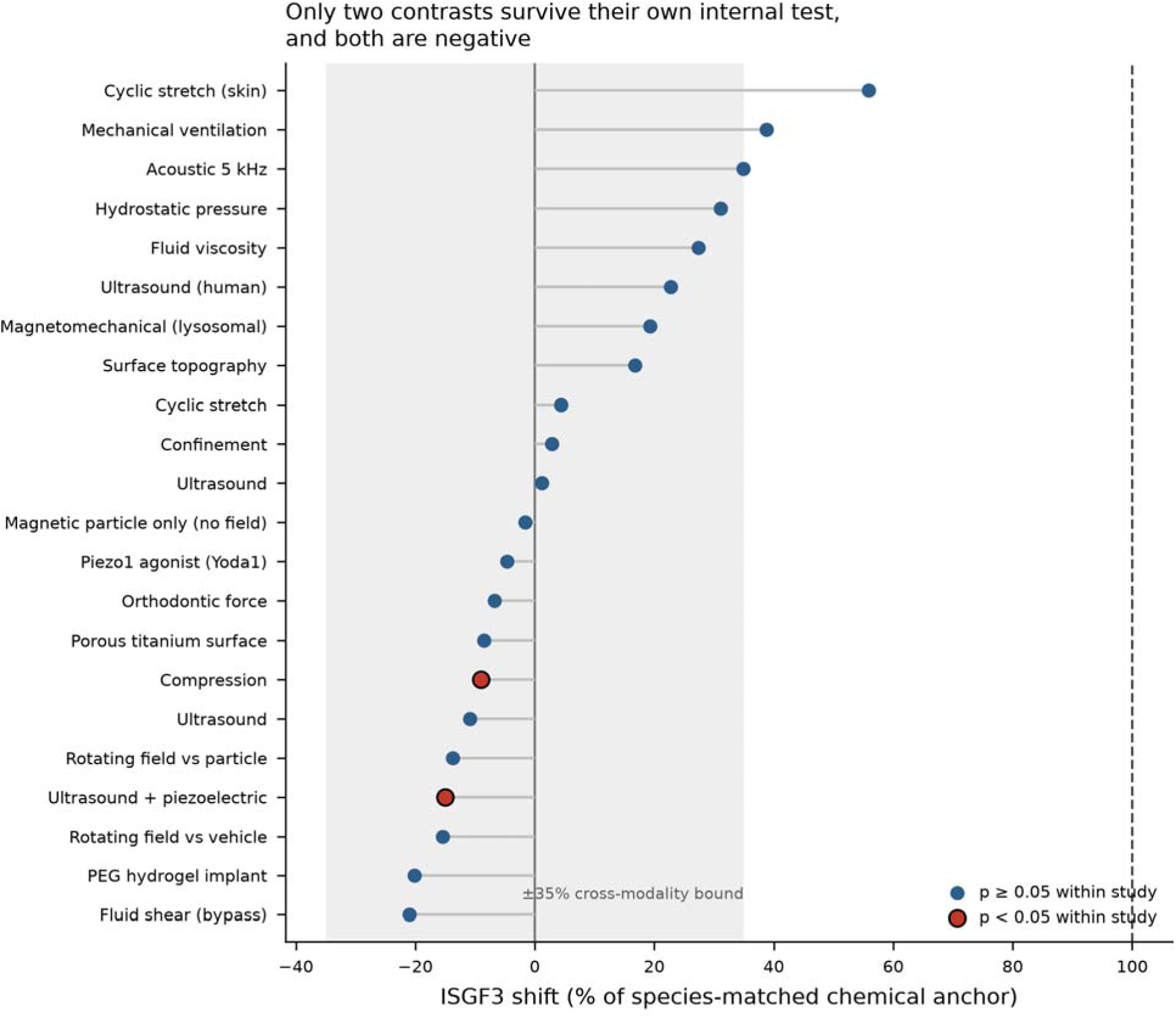
Modality projection. ISGF3 shift as a percentage of the species-matched chemical anchor for 22 contrasts across ten mechanical modalities, grouped by modality family (n per family at left), with the Piezo1 agonist shown separately as a pharmacological surrogate. Red markers are the two contrasts that reach within-study permutation p < 0.05: neither exceeds 15% of the anchor, and both are negative. Every shift beyond 20% of the anchor comes from a design whose own permutation test returns p ≥ 0.10.

The decisive qualification is which of these survive within-study inference. Only two contrasts reach within-study permutation p < 0.05, and neither exceeds 15% of the chemical anchor: ultrasound on a piezoelectric membrane (−15.0%) and compression (−9.0%) (Figure 7). Both are negative. Every shift larger than 20% of the anchor, in either direction, comes from a design whose own permutation test returns p ≥ 0.10. We report a two-tier bound: within the stiffness family the equivalence margin is ±10%. Across all ten modalities 20 of the 22 contrasts fall within ±35% of the anchor, and ±35% is the defensible cross-modality bound.

It is worth walking the modality table in the direction the data imposes rather than the direction a mechanism would suggest: the ordering is informative in itself. The largest absolute shift in the entire projection, in-vivo cyclic stretch of skin (+55.9% of the anchor, GSE125806), comes from whole skin rather than sorted myeloid cells. Composition and macrophage-intrinsic response are confounded here exactly as they are in the two whole-tumor studies demoted from the main pool, and its within-study permutation p is 1.0, the maximum the design can return. The second largest, mechanical ventilation (+38.8%, GSE274828, lung myeloid cells pseudobulked from single-cell data), returns p = 0.50 on a design with only four possible label assignments. The third, cyclic hydrostatic pressure on *Piezo1*-floxed BMDM (+31.1%, GSE133069), is a one-versus-one comparison and likewise returns p = 1.0. At the negative extreme, in-vivo fluid shear during cardiopulmonary bypass (−21.0%, GSE143780) is a five-donor paired design whose attainable floor is 2/32 = 0.063 and which returns p = 0.125. PEG hydrogel implantation (−20.2%, GSE185761) is read from foreign-body-response capsule macrophages recovered at day 14, a setting in which the comparator is “no implant” and so differs in tissue state as well as in mechanics (p = 0.40). The two contrasts that do survive within-study inference are both smaller than the median absolute shift of 15.2%: low-intensity ultrasound delivered through a piezoelectric membrane (−15.0%, p = 0.029) and compression of RAW264.7 in agarose at 0.14 kPa (−9.0%, p = 0.004). No contrast in this corpus both moves the module by more than a fifth of the chemical scale and passes its own internal test. Both survivors are negative. Where mechanics does move this module detectably, it moves it down.

Pooling within modality families was attempted, but acoustics/ultrasound (K = 5) and topography/materials (K = 3) have I² = 92.4% and 86.4%. These are real directional disagreements: the pooled values (+5.5% and −1.3% of the anchor) are not interpretable as modality effects.

### 3.6 The YAP/TAZ arm: an anchor exists, but the module is not reliable enough to carry it

A mechanistic account in which stiffness acts on ISGs through YAP/TAZ requires, at minimum, that the YAP module move under mechanical load. Screening all YAP contrasts for within-study permutation p < 0.05 and random-set p < 0.05 yields a single anchor: porous Ti vs plastic (GSE99445, 8 paired donors, Δ = +0.01085, p_perm = 0.039, p_rand = 0.035, 40 DE genes). The nearest miss is Yoda1 vs DMSO (E-MTAB-15203, human MDM, 6 paired donors, Δ = +0.01707, 4,421 DE genes), which clears neither test: its permutation p of 0.063 is the second attainable level for a six-donor paired design, and its random-set p is 0.108. We report margins against both, so that the bound does not rest on a single contrast. A compression contrast (GSE304867) was rejected on both tests (p_perm = 0.303, p_rand = 0.641), and its YAP shift is negative rather than positive.

The stiffness family shows no YAP movement against these anchors: pooled μ = −0.00052, 95% CI [−0.00267, +0.00164] (K = 10, I² = 0.0%) (Figure 6B). This is equivalent to zero at ±20% of the Yoda1 near-miss value (p = 0.0070) and at ±50% of the porous-Ti anchor (p = 3.0 × 10□□), though not at ±20% of the porous-Ti anchor (p = 0.058). Taking the qualifying anchor as the ruler, the conservative reading is that the stiffness family is equivalent to zero on the YAP module at a ±50% margin, not ±20%.

Two findings argue against building the mechanistic claim on this module. First, the qualifying anchor and the near-miss differ by 1.57-fold, and two measurements of the same chemical stimulus in two sub-series of one cohort differ by 2.44-fold (+0.02684 vs +0.01099) (Figure 3B). Second, the YAP module is the only one of the four to show a detectable small-study effect (Egger p = 0.033) (Figure 6B). We report the YAP result as a bounded absence with an explicit factor-2.4 scale uncertainty and do not use it to adjudicate the upstream mechanism.

A species correction follows from the same anchors. The frequently repeated statement that chemical M1 does not engage YAP holds only in mouse (chemical anchor 0.00343); the human chemical anchor is 0.0110–0.0268 (median 0.0189), the same order as the Yoda1 mechanical anchor. Any such claim must be made per species.

### 3.7 Placement of a magnetic actuator does not change the ISG answer

A three-arm in-vivo dataset (GSE275018; 94,218 cells, 18,244 myeloid cells pseudobulked) separates the particle from the field (Figure 8). Iron loading alone (nanomotor vs PBS) gives Δ_ISGF3 = −0.0021 with zero genes at FDR < 0.05; the rotating field on top of loading (RMF vs nanomotor) gives −0.0171 with 780 DE genes; the combined contrast gives −0.0192 with 1,176 DE genes (Figure 8B). The entire transcriptional effect belongs to the field, not the particle. Its direction is downward. In a second magnetomechanical dataset (GSE273977) an apparent +0.0240 recovers only 29.7% of an IL-4-imposed suppression of 0.0807, measured on the same array platform in GSE69607, where polarization to IL-4 M2 lowers the module from 0.2987 to 0.2180, and ends 0.0567 below the M0 baseline: a partial release from a floor, not induction. GSE273977 also lacks a particle-only arm, so iron loading and field actuation remain confounded within it.

**Figure 8.**
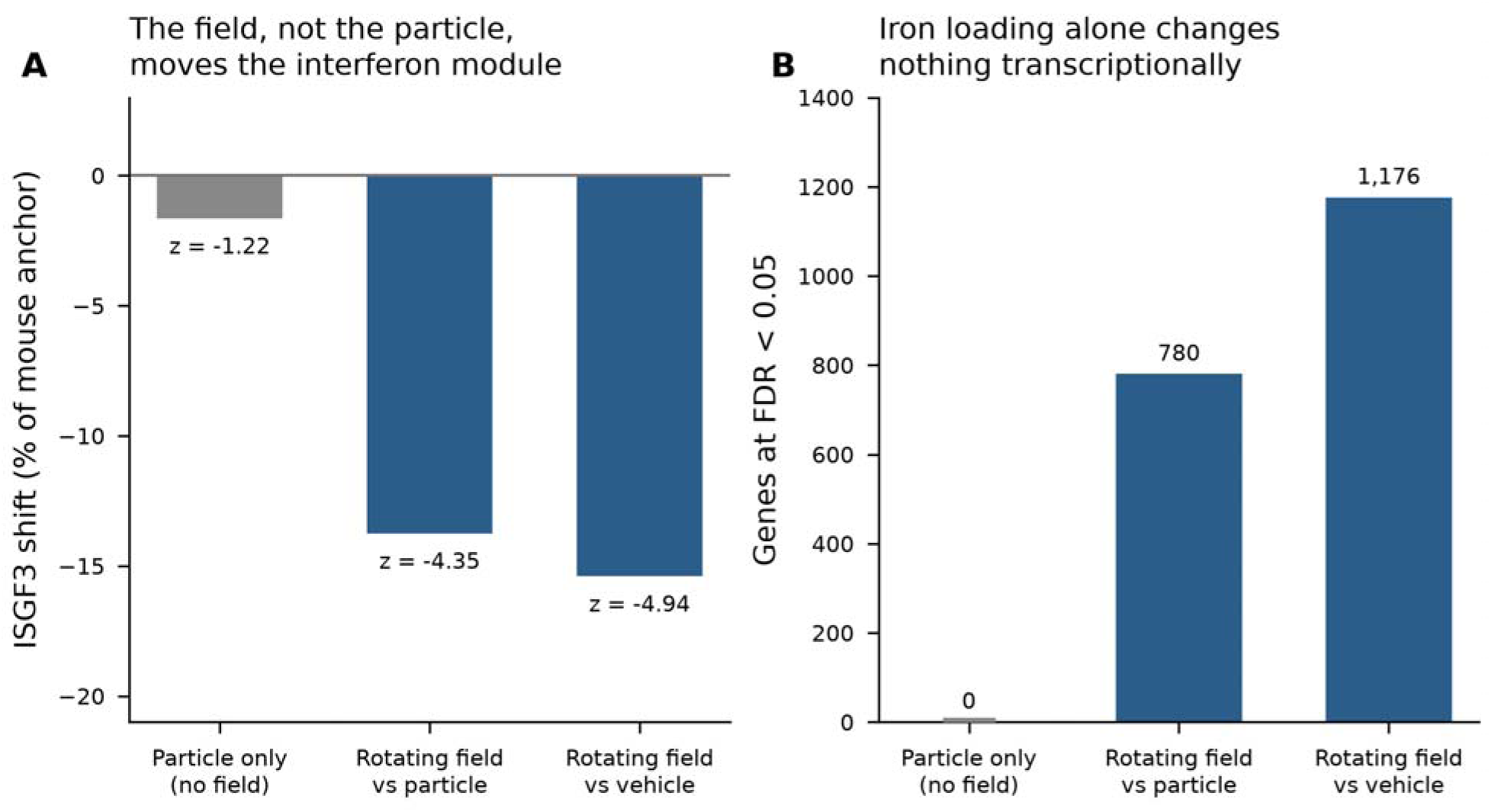
A three-arm in-vivo magnetomechanical experiment separates the particle from the field (GSE275018, 18,244 myeloid cells pseudobulked). (A) ISGF3 shift as a percentage of the mous chemical anchor, with the random-set z statistic. (B) Genes differentially expressed at FDR < 0.05 in th same three contrasts. Iron loading alone changes nothing. The entire transcriptional effect appears when the rotating field is applied, and its direction is downward.

Interferon-stimulated genes are not a readout that can discriminate where a mechanical actuator is placed. Scoring the same three arms on the effector module shows what the field does instead (§3.9).

### 3.8 The one dataset with a true dose gradient trends upward

Every other dataset in the corpus compares two mechanical levels. One does not: E-MTAB-7900 profile rat bone-marrow-derived macrophages on substrates of 0.1, 1, 10 and 100 kPa in four donors, giving the only four-point gradient available (Figure 9A). Fitting the module score against log10 stiffness gives a slope of +0.0142 singscore units per decade (SE 0.0062, t = 2.27, p_trend = 0.108, Spearman ρ = 0.558) (Figure 9B). The two Hallmark modules trend the same way and at a similar magnitude: Hallmark IFN-α +0.0125 per decade (p = 0.120, ρ = 0.473) and Hallmark IFN-γ +0.0106 per decade (p = 0.120, ρ = 0.485). The YAP/TAZ module shows no trend at all (−0.0006 per decade, p = 0.944, ρ = −0.133). Expressed on the chemical scale, the ISGF3 slope is +11.4% of the human anchor per decade of stiffness. Reaching the chemical anchor along this line would require roughly nine decades of stiffness, a range that does not exist in tissue.

**Figure 9.**
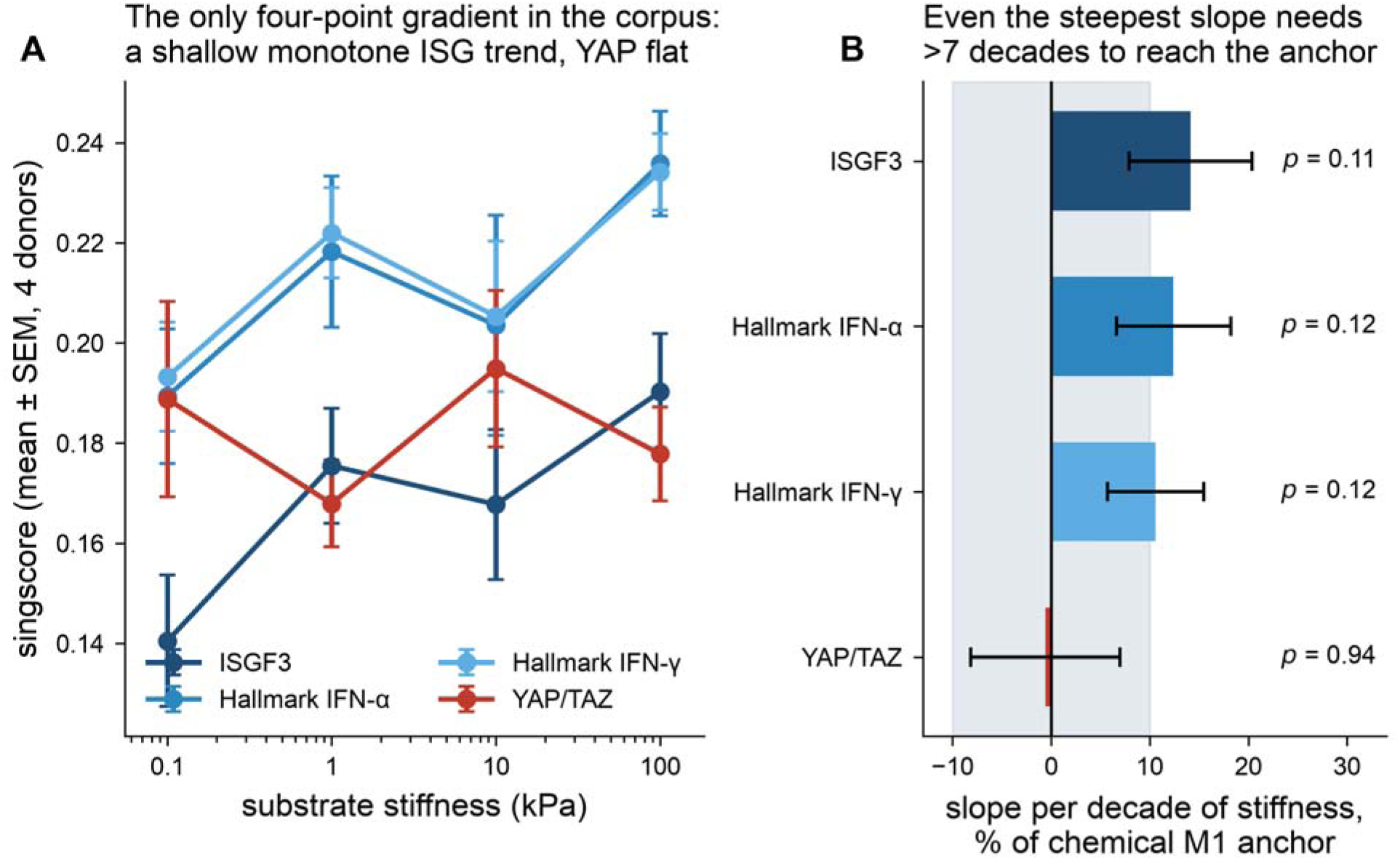
The only four-point dose gradient in the corpus. (A) Module scores for rat BMDM across 0.1, 1, 10 and 100 kPa (E-MTAB-7900, 4 donors, mean ± SEM). Three ISG definitions trend upward together, while the YAP/TAZ module does not. (B) Slope per decade of stiffness expressed as a percentage of the chemical M1 anchor, with standard errors and trend-test p values. The steepest slope, +11.4% of the anchor per decade, would require roughly nine decades of stiffness to reach the chemical response.

Three features of this result deserve separate treatment. First, it is not significant: with four donors th trend test returns p = 0.108, and the slope’s confidence interval includes zero, so it cannot overturn the pooled estimate. Second, it is nonetheless internally coherent. Three ISG definitions trend together while the mechanotransduction module does not (Figure 9A), a pattern that is not what a single noisy donor driving the fit would produce. Third, its endpoint contrast (0.1 vs 100 kPa, delta = +0.0498, +40.0% of the human anchor) is the largest positive study-level value in the main pool and the only one whose 95% confidence interval excludes zero on the positive side ([+0.0026, +0.0970]). The leave-one-out analysis shows that removing this study moves the pooled estimate only from −0.89% to −0.60% (Figure 6A): it is not driving the overall result. It is, instead, the single observation in the corpus most likely to be a real effect that the pooled design is not built to detect.

The asymmetry between this dataset and the rest is a design asymmetry rather than a biological one, worth stating precisely. A two-level comparison at 1 kPa versus 50 kPa samples a putative dose-response curve at two points and reports the chord between them. If the true relation is shallow and monotone, as the slope above would imply, a chord drawn across the physiological range is small and will be reported as “no effect.” The pooled estimate of −0.9% is best read as a statement about chords, the +11.4%-per-decade slope as a statement about the only gradient available. The two are not in conflict. Resolving which describes the biology requires gradient designs, not more two-level ones.

### 3.9 The effector arm moves where the interferon arm does not

Every contrast above was rescored on the NF-κB effector modules: same expression matrices, same unfiltered background, same pipeline. The two arms differ only in which genes are read. One background is used throughout (§2.4), so the percentages in this section are directly comparable with those of §3.2–§3.8.

Under static stiffness, neither arm moves (Figure 10B). Pooling the eleven stiffness and applied-force contrasts for which both arms could be scored on identical matrices gives ISGF3 +2.80% of anchor (95% CI −5.67 to +11.27, I² = 78.2%), TNFA/NF-κB −5.27% (−11.36 to +0.82, I² = 54.3%), inflammatory response −3.15% (−11.87 to +5.58, I² = 89.7%) and the field panel −5.88% (−13.43 to +1.67, I² = 84.4%). All four are statistically equivalent to zero at a ±20% margin (TOST p ≤ 1.0 × 10□³). At the tighter ±10% margin the interferon arm is still equivalent (p = 0.044); none of the three effector modules is (p = 0.057, 0.055 and 0.126). This is not because the effector estimates are larger: it is because they are more dispersed across studies. The paired within-study separation, effector percentage minus interferon percentage, is −5.5 points (SD 31.3, t p = 0.572, Wilcoxon p = 0.465) (Figure 10B). The two arms are not merely both null on average. They are uncorrelated across studies (Pearson r = −0.14) and point in opposite directions in 6 of 11.

**Figure 10.**
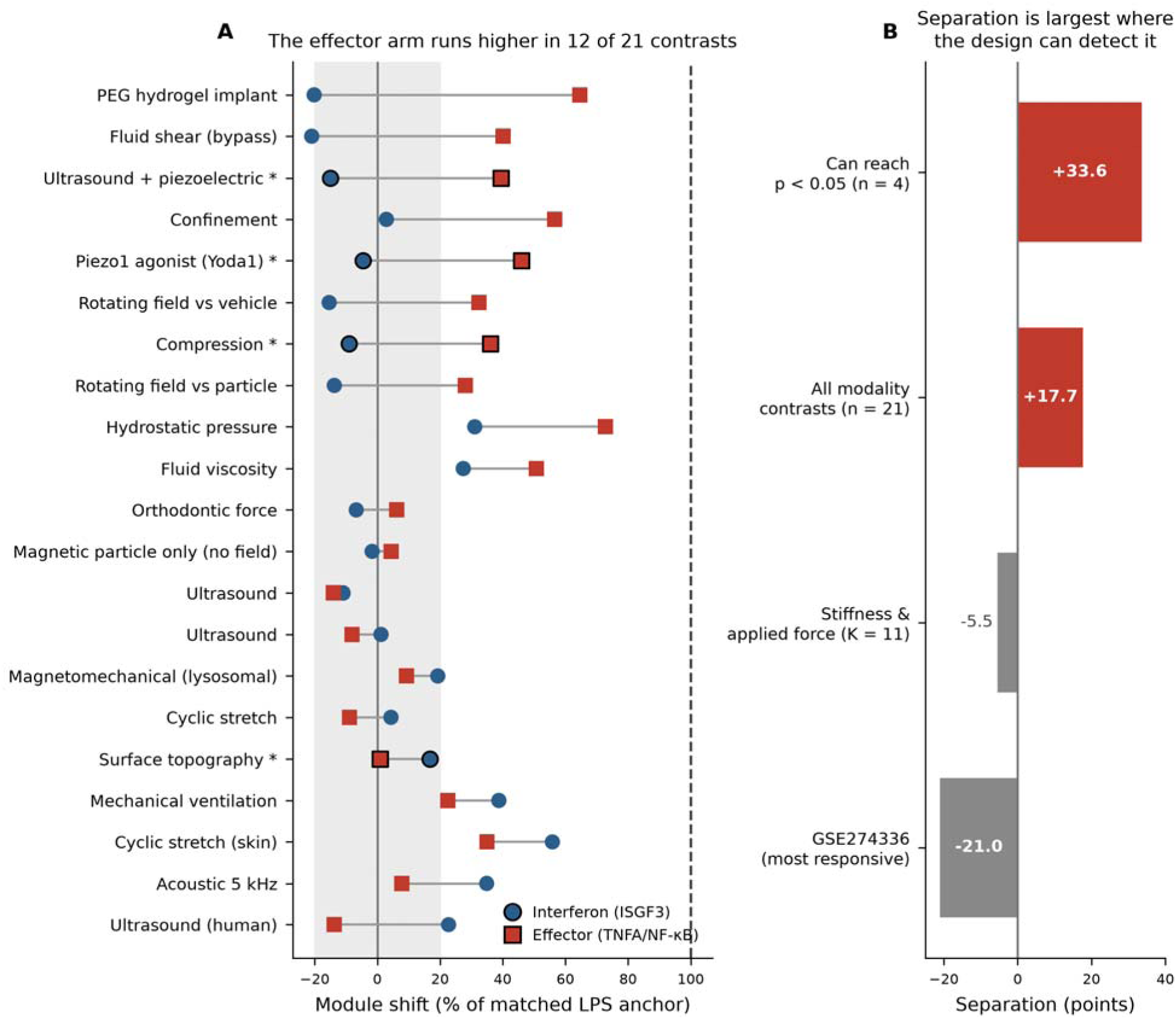
The two arms across all mechanical modalities. (A) Each row is one contrast; blue circles are the ISGF3-specific interferon module and red squares the TNFA/NF-κB effector module, both expressed as a percentage of the matched LPS anchor and joined by a grey connector. Black outlines mark within-study permutation p < 0.05; an asterisk on the row label marks the four designs whose permutation space is large enough to return p < 0.05 at all. The shaded band is the ±20% equivalence region, and the dashed line is the anchor. Both arms vary widely, and the effector arm runs higher in 12 of the 21 contrasts. (B) Mean separation (effector percentage minus interferon percentage) for four groupings: the eleven stiffness and applied-force contrasts (−5.5 points), all 21 modality contrasts (+17.7), the four contrasts whose designs can reach significance (+33.6), and the single most transcriptionally responsive contrast in the corpus. The separation is largest where the design can detect it, and its sign is not fixed.

This null is interpretable, not vacuous: the effector arm had room to move. Absolute effector scores in the control arms of these eleven contrasts span 0.096 to 0.405. The chemical anchors start from 0.219 in mouse and 0.352 in human, and LPS then adds +0.098 and +0.084, carrying them to 0.318 and 0.436. Every one of the eleven mechanical contrasts begins below the level the chemical anchor demonstrably reached in the same species. A ceiling cannot explain the absence of a shift.

Where mechanical loading registers, the arms separate (Figure 10). The modality projection of §3.5 was rescored on both arms. One of its 22 contrasts was excluded: the porous-titanium arm of GSE99445, which is a mechanical anchor for the YAP module (§2.6) and so is not independent of the anchor set. That leaves 21 contrasts: 16 bulk and 5 obtained by myeloid pseudobulk from single-cell data. Across them the effector arm sits a mean of +17.7 percentage points above the interferon arm (SD 35.2, median +12.9, positive in 12 of 21, paired t p = 0.0322, Wilcoxon p = 0.0595; Figure 10A). The interferon arm averages +6.5% of its anchor and the effector arm +24.2%. Across contrasts the two are uncorrelated (Pearson r = −0.05, p = 0.82). The separation is not a common scaling of both arms by a per-study factor, which would have produced a positive correlation.

The design limitation of §3.5 applies with full force here: only 4 of the 21 contrasts have a permutation space large enough to return p < 0.05 at all. Within those four, the effector arm is significant in three and the interferon arm in two. The informative cases are the two in which both arms clear significance: there, the arms move in opposite directions (Figure 10A).

- compression of bone-marrow-derived macrophages (GSE304867; permutation space 924): interferon **−9.0%** (p = 0.004), effector **+36.0%** (p = 0.002);
- ultrasound on a piezoelectric substrate (GSE236912; space 70): interferon **−15.0%** (p = 0.029), effector +39.5% (p = 0.029), both at the design floor of 2/70.

In the third resolvable design only the effector arm moves: the Piezo1 agonist Yoda1 in human monocyte-derived macrophages (E-MTAB-15203; six paired donors, permutation space 2 = 64) gives interferon −4.6% at p = 0.063, the second attainable level, against effector +46.0% at p = 0.031, the floor (Figure 10A). The fourth design, nano/microstructured sulfonated PEEK topography (GSE290857; space 70), moves neither arm decisively (interferon +16.8%, p = 0.057; effector +0.9%, p = 0.83). It is the one design in this subset where the interferon arm is the larger of the two. Averaged over the three contrasts in which the effector arm is supported by within-study inference, the effector arm reaches +40.5% of its anchor while the interferon arm of the same contrasts sits at −9.5%: a separation of 50 points. Figure 10B places these alongside the stiffness pool. The separation is −5.5 points where neither arm moves, +17.7 points across all modalities, and +33.6 points in the four designs that can resolve it.

The direction of the effector shift is not fixed. The single most transcriptionally responsive contrast in the corpus is the one used in §3.2 to show that the interferon null is not a sensitivity failure (GSE274336, 12 vs 12, stratified by harvest time, 4,318 genes at FDR < 0.05) (Figure 5). Rescored on both arms against 1,000 size-matched random gene sets drawn from the same 33,043-gene background, it reproduces the dissociation with the largest margin anywhere in this work, and the effector arm points down: ISGF3 Δ = −0.00254 (random-set z = −0.94, p_random = 0.509), TNFA/NF-κB Δ = −0.02243 (z = −8.04, p_random < 0.001), inflammatory response −0.01287 (z = −4.74), field panel −0.07301 (z = −4.83, p_random = 0.001), YAP/TAZ +0.00049 (z = −0.10, p_random = 0.954). On one set of samples, one pipeline and one null model, the interferon module sits in the middle of its random-set distribution. The effector module sits eight standard deviations outside it (Supplementary Table S15). The effector arm here moves downward, opposite in sign to the modality mean (Figure 10B). That is the point: the two arms are dissociable, not reciprocally coupled in a fixed direction.

The magnetomechanical three-arm experiment separates particle from field on both arms (Figure 8). Rescoring GSE275018 (§3.7) gives, for iron loading alone, interferon −1.6% (random-set z = −1.22) (Figure 8A) and effector +4.4% (z = −0.40) (Figure 10A). Neither arm moves. Adding the rotating field moves both, in opposite directions: interferon −13.8% (z = −4.35) and effector +28.0% (z = +2.18); the combined contrast gives −15.4% (z = −4.94) and +32.4% (z = +1.67). Permutation p for the field arms is 0.1, the attainable floor for this 2 × 10 design, so the within-study test cannot certify either arm on its own. What separates them is the random-set statistic, and it does so in opposite directions. A rotating magnetic field does not leave tissue macrophages unchanged. It redistributes them: the effector arm up, the interferon arm down.

### Source tables

Every quantity reported above is reproduced in the Supplementary Tables: study-level effect sizes for the main pool (Table S4), leave-one-study-out refits (Table S5), species strata (Table S6), module-level results with Egger and Begg tests (Table S7), the Monte-Carlo power curve (Table S8), the chemical reference anchors (Table S9), the complete modality-projection scoring table (Table S10), and the four-module effect sizes for the expanded K = 12 pool (Table S11). The two-arm quantities of §3.9 appear across four further tables: the effector-ruler validation (Table S12), the study-level two-arm effects for the eleven stiffness and applied-force contrasts (Table S13), the two-arm projection across all 21 modality contrasts with each design’s attainable minimum p (Table S14), and the five-module result for GSE274336 with its detected gene background (Table S15).

## 4 Discussion

### 4.1 What the result is, stated narrowly

Across ten independent transcriptomic studies, mechanical perturbation of macrophages shifts the ISGF3-dependent interferon-stimulated gene module by −0.9% of the shift produced by LPS + IFN-γ, with a 95% confidence interval that excludes anything larger than ±6.2%. The claim we are entitled to make is not “mechanics does nothing,” nor is it “mechanics suppresses interferon signaling.” It is narrower, and more useful: *whatever mechanically induced “M1-like” states are, they are not the chemical M1 program as seen through its interferon arm*.

The narrowness matters: two of our own analyses forbid the stronger readings. The pooled sign flips positive when a single study is removed (§3.4), so direction is not established. The modality projection, meanwhile, shows mechanics is demonstrably not inert toward ISGs. It moves them in both directions depending on modality (§3.5). What remains defensible is a statement about magnitude relative to a chemical reference. That is why the analysis was built around an anchor instead of a null hypothesis.

### 4.2 Why the field reported “stiffness drives M1”, and why that is not contradicted

The mechanoimmunology literature contains apparently irreconcilable reports: stiff substrates have been described as driving M1 (4,6) (the latter TRPV4-dependent), soft substrates as driving M1 (7), and stiffness as pushing tumor-associated macrophages toward M2 (8). Our result does not adjudicate that dispute: it explains why it can persist.

Table 2 sets the principal reports side by side with the readout each of them used, and the pattern is immediate: the disagreement is not between laboratories measuring the same thing but between laboratories measuring different arms of the program, with different instruments. Of the seven stiffness-direction studies that recur in the field’s contradiction tables, five read NF-κB/AP-1 effector output by qPCR, ELISA or flow cytometry, one reads tissue-level histology, and one reads single-cell composition in a tumor. None reports the interferon arm. A reader who assembles these into a single column headed “effect of stiffness on M1” is, in effect, averaging a cytokine secretion assay, a capsule-thickness measurement and a cell-type proportion. The average should not be expected to have a stable sign.

**Table 2.** Principal stiffness-direction reports, the readout each used, and the arm of the M1 program that readout samples.

| **Study** | **System** | **Reported direction** | **Primary readout** | **Arm of M1 sampled** |
|---|---|---|---|---|
| Blakney et al. (6) | PEG hydrogels, in vitro and in vivo | stiffer → more inflammatory | cell density, capsule thickness, histology | tissue-level host response |
| Dutta et al. (4) | TRPV4, PDMS of graded stiffness | stiffer → M1 | iNOS, TNF-α, IL-6 (qPCR, protein) | NF-κB / AP-1 effector |
| Chen et al. (7) | BMDM on polyacrylamide | softer → M1, via NF-κB | iNOS, TNF-α, IL-1β | NF-κB / AP-1 effector |
| Guenther (8) | DC/macrophage co-culture, graded stiffness | stiffer → tumor-associated (M2-like) | surface markers, secreted cytokines | surface phenotype + effector |
| Patel et al. (17) | macrophage cell elasticity | elasticity determines function | phagocytosis, migration, spreading | cellular function |
| Friedemann et al. (45) | 3D collagen, 27.1–118.5 Pa, GAG-functionalized | stiffness does not track output; GAG does | IL-10, IL-12, TNF-α secretion | NF-κB / AP-1 effector |
| Taufalele et al. (18) | mammary tumor stiffness in vivo | stiffer → M2-like accumulation | single-cell composition | myeloid composition |

Notes: The classification in the final column is ours; the "reported direction" column reproduces each study's own conclusion. No study in this set reports the ISGF3-dependent interferon arm, the quantity measured here.

Nearly all of those studies read out iNOS, TNF-α, IL-6, IL-12, CD80 by qPCR, flow cytometry or ELISA. Those are NF-κB and AP-1 effectors: part of the M1 phenotype, but not the interferon arm, and nothing in our analysis says they are unmoved by mechanics. What our analysis says is that the ISG module, the part of chemical M1 most specific to it, does not come along. A mechanically induced state that raises TNF-α without raising ISGs is a different transcriptional state from LPS + IFN-γ, even if both are labelled “M1”. The field’s sign disagreements are not evidence of irreproducibility. They are evidence that “M1” is being used for a set of states that a single axis cannot order.

The Friedemann entry in Table 2 deserves expansion. We initially mis-filed it as a stiffness-direction result, and the correction is instructive. Friedemann et al. (45) functionalized 3D collagen networks spanning 27.1–118.5 Pa, two to three orders of magnitude softer than the rest of the stiffness literature cited here, and found that EDC crosslinking (57.5 Pa) shifted macrophages toward an anti-inflammatory profile while the stiffer HA (73.6 Pa) and sHA (118.5 Pa) networks did not. The authors state explicitly that cytokine output does not track network stiffness. The result is real. But it is evidence that glycosaminoglycan presentation can override a stiffness effect within a single stiffness range, not that stiffness has a direction. Several entries in the field’s “contradiction” tables may be of this kind.

Two further details of that study sharpen the point. Its stiffness values were obtained by colloidal-probe atomic force microscopy with a Hertzian fit, so they are effective moduli of a hydrated fibrillar network, not commensurable with the bulk moduli of polyacrylamide or PDMS substrates quoted elsewhere in Table 2. A 100 Pa collagen network and a 100 Pa polyacrylamide gel do not present the same mechanical environment to a cell that must deform individual fibres. And every arm of the experiment received LPS in the final 24 h, with no LPS-free control. The study could not have entered our corpus under criterion 3 even if its stiffness range had overlapped the rest. A field-level contradiction table that places this study in the same column as a 1–50 kPa polyacrylamide experiment is comparing neither the same stimulus nor the same baseline.

### 4.3 Mechanics is bidirectional toward ISGs, and that is the second finding

Projecting the same score across ten non-stiffness modalities gives shifts from −0.0251 to +0.0696 that separate cleanly. In-vivo fluid shear, implanted hydrogel, a rotating magnetic field, ultrasound-on-piezoelectric-membrane and compression lower the module. Cyclic stretch, mechanical ventilation, acoustic oscillation, hydrostatic pressure and fluid viscosity raise it (Figure 7).

We resist reading mechanistic meaning into that split for one reason: only two of the 22 contrasts survive within-study permutation, and neither exceeds 15% of the chemical anchor (Figure 7). Everything larger comes from contrasts their own internal tests do not support. Pooling within families does not help either: acoustics/ultrasound and topography/materials carry I² of 92.4% and 86.4%, genuine directional disagreement rather than noise around a common effect.

The honest summary is a two-tier bound: ±10% within the stiffness family, ≈±35% across all mechanical modalities.

### 4.4 Amplitude does not buy interferon engagement

A natural objection to an equivalence result is that the mechanical perturbations were simply too weak. Figure 5 addresses that directly, plotting each contrast’s ISG shift against the number of genes it moves at FDR < 0.05: a crude but assumption-free measure of how much transcription the perturbation actually changed. Across the 20 mechanical contrasts for which both a differential-expression count and an anchor percentage are available, the shifts span −21.0% to +38.8% of the anchor, 12 of the 20 fall outside the ±10% band, and the absolute size rises with amplitude only weakly, without reaching significance (Spearman ρ = 0.38, p = 0.097). The picture is not one of mechanical perturbations that do nothing. Amplitude and ISG displacement are at most loosely coupled, as one would expect if large transcriptional rearrangements occasionally graze the module.

What the figure shows instead is a ceiling. No mechanical contrast reaches the chemical anchor at any amplitude. The highest-amplitude mechanical contrasts in the corpus move 5,000–7,300 genes and still sit at or below 27% of the anchor; the chemical anchors move 2,459 and 3,631 genes and sit, by construction, at 100%. The clearest single case is GSE274336: 4,318 genes differentially expressed, a −2.0% ISG shift. Chemical M1 and mechanical activation are not distinguished by how much transcription they change (they are comparable on that axis) but by *which* transcription they change. An objection based on insufficient perturbation strength would need a mechanical contrast that both moves thousands of genes and moves the module. The corpus contains the first without the second.

### 4.5 One genuine tension we cannot resolve

The only dataset in the corpus with a true dose gradient, rat BMDM across 0.1, 1, 10 and 100 kPa (E-MTAB-7900) (Figure 9), trends upward: ISGF3 rises by +0.0142 per decade of stiffness (SE 0.0062, p_trend = 0.108, Spearman ρ = 0.558), with Hallmark IFN-α and IFN-γ trending the same way (+0.0125 and +0.0106, both p = 0.120). YAP shows no trend (p = 0.944). The endpoint contrast here (+0.0498, +40% of the anchor) is the largest positive value in the pooled set.

With four donors the confidence interval crosses zero and the trend is not significant. It does not overturn the pooled result, but it is the single observation most likely to be true and missed. Almost every dataset in this field is two-level, and a field that tests stiffness at two levels and finds nothing may simply be sampling a dose-response curve at two points. This is the highest-value experiment a reader could run, not a caveat to be filed away.

### 4.6 The YAP/TAZ route cannot be adjudicated with current modules

A natural mechanistic hypothesis is that stiffness reaches ISGs through YAP/TAZ, which is reported to restrain TBK1 (5). We can now test the premise: mechanical YAP movement is detectable on porous titanium topography (+0.0109), the one contrast that clears the anchor screen, and short of that threshold in Yoda1 in human MDM (+0.0171). Against those anchors the stiffness family shows no YAP movement at all (μ = −0.00052, 95% CI [−0.00267, +0.00164], I² = 0.0%).

We nonetheless decline to call the route excluded, for two measurement reasons. The qualifying anchor and the near-miss differ 1.57-fold, and two measurements of the same chemical stimulus in two sub-series of one donor cohort differ 2.44-fold (+0.0268 vs +0.0110), while the ISG module in those same sub-series agrees to 1.5% (Figure 3B). The Wang 22-gene signature is not a reliable ruler at the precision this question needs. YAP is also the only module of four showing a detectable small-study effect (Egger p = 0.033) (Figure 6B).

One correction belongs in the literature regardless. The statement that chemical M1 does not engage YAP is mouse-specific: the mouse chemical anchor is 0.0034, but the human chemical anchor is 0.0110–0.0268, the same order as a Piezo1 agonist. Any cross-species argument built on “chemical polarization leaves YAP alone” needs restating.

### 4.7 Where a magnetic actuator sits does not change the ISG answer

A three-arm in-vivo experiment lets the particle be separated from the field (Figure 8). Iron loading without actuation produces zero differentially expressed genes and Δ_ISGF3 = −0.0021; adding the rotating field produces 780 genes and −0.0171. The transcriptional effect belongs entirely to the field, and it points downward.

Reading the same three arms on the effector module changes the picture from an absence to a redistribution. Iron loading alone leaves both arms where they were (interferon −1.6%, effector +4.4%). The rotating field moves both, in opposite directions: the effector arm up by 28.0% of anchor and the interferon arm down by 13.8%, each outside its size-matched random distribution (z = +2.18 and −4.35). So a rotating field does not do “nothing” to a tissue macrophage, and it does not deliver “M1 repolarization”. It shifts one axis of the inflammatory program up and another down.

This has a direct consequence for magnetomechanical therapeutics, where “M1 repolarization” is a common stated goal and interferon-related readouts are often used to support it. On this evidence, ISGs are the wrong endpoint for discriminating lysosomal from membrane-anchored actuation: both placements are silent in that dimension, and the effector arm responds to the field rather than to the particle. Neither readout distinguishes placement on its own. Endpoints that could discriminate are those specific to the proposed lesion: galectin-3/8 recruitment and lysosomal membrane permeabilization, TFEB translocation and lysosomal biogenesis, and AMPK–ULK1 signalling. The iron-only arm, with zero differentially expressed genes, is an unusually clean negative control for exactly that experiment.

### 4.8 The two arms come apart, and that is the shape of the result

The question this paper opened with (is mechanically induced “M1” the chemical M1 program?) can now be answered with a structure rather than a single number. The chemical program has at least two separable arms. Mechanical input does not treat them alike.

Three independent lines converge on that reading, and they are not the same experiment repeated. The first is a within-contrast dissociation in the corpus’s most responsive dataset: the interferon module sits inside its random-set distribution, while the effector module sits eight standard deviations outside it, on the same samples. The second is the modality projection. Here the arms separate by a mean of 17.7 points, and in the two contrasts powerful enough for both arms to clear significance, they move in opposite directions. The third is the in-vivo magnetomechanical experiment, where the arms separate by 48 points under the field and by 6 points under the particle alone. These differ in species, platform, the physical input applied, and the inferential machinery used. That is why their agreement carries weight.

Two features of that structure resist a simple slogan, and both are properties of the data rather than of our caution. The sign of the effector shift is not fixed: compression, ultrasound, Piezo1 agonism and a rotating magnetic field all raise it, while the stiffness contrast of GSE274336 lowers it by 23 points. Under static substrate stiffness the arms do not separate at all: the pooled effector estimate is −5.3% of anchor with a confidence interval that excludes a 20% effect, measured in studies whose control arms sit below the anchor’s own controls and had room to move. “Mechanics activates the effector arm” would be the wrong summary. What the data support is narrower and more useful: where a mechanical input registers transcriptionally at all, it registers on the effector arm and not on the interferon arm, in whichever direction it registers.

This also disposes of a reading of our main result that a corpus selected for LPS-free arms might be a corpus of weak perturbations, in which any module would fail to move. We raised this possibility ourselves in §4.9. It is now testable rather than arguable: in the same studies, with the same matrices, one module fails to move and another moves, downward in GSE274336, upward under compression and ultrasound. A corpus too inert to register would not produce z = −8.0.

Finally, the dissociation is not an artefact of module overlap or of module construction. The two sets share 5 of 82 human genes, so they are close to independent populations. One asymmetry does remain: occupancy-defined interferon module against expression-defined effector modules (§2.3). But this works against the dissociation rather than for it. An expression-derived set is the more permissive of the two, and would be expected to move *more* readily in a corpus of inert perturbations, not less.

### 4.9 Limitations

The two arms are not measured with equal authority. The interferon module is defined by measured ISGF3 promoter occupancy; the effector modules are expression-derived Hallmark sets, because no RELA occupancy set of comparable construction is distributed (§2.3). A reader should treat “the interferon arm did not move” as the better-supported of the two statements, and “the effector arm moved” as the one resting on a more permissive module definition. The human effector percentages also carry a between-cohort uncertainty: roughly 10% for the two large modules and 22% for the six-gene panel, read off the two independent human LPS cohorts (§2.6). Absolute human effector magnitudes are reported, but at that precision rather than better.

Concluding that mechanics fails to activate macrophages inverts the argument. The modality projection contains mechanical contrasts that move thousands of genes, and §3.9 shows which arm they move. The claim is about routing, not magnitude.

The human evidence base is three studies. A study-level permutation over three studies cannot return a p below 0.25: the human equivalence statement (±20%, p = 0.032) rests on a parametric test with two degrees of freedom and should be read as directional. Four to six human studies would change this, and the obstacle is availability rather than feasibility. The one human monocyte shear dataset we identified (GSE291575) returns HTTP 403 for all supplementary files. This is an archive-availability bias specific to repository-based meta-analysis, and no standard publication-bias test addresses it.

Ten studies are not ten independent units. The Kish effective number under random-effects weights is 5.72. Statements about breadth of replication should use that figure.

The equivalence margin is post hoc. We set it by what the design can resolve, an MDE at 80% power of 8.8% of the anchor, not by a biological threshold. For that reason, we report the estimate as the primary quantity.

Small-study tests are weak here and should not reassure. Egger and Begg are null for all three ISG modules, but at K = 10 they have little power: the raw Δ-versus-SE correlation is −0.58, and the least precise studies sit on the negative side. That is the same instability that flips the pooled sign under leave-one-out.

Two in-vivo whole-tumor studies were excluded from the main pool because bulk tissue cannot separate myeloid composition from macrophage-intrinsic response. Including them changes the estimate by less than 0.01 percentage points of the anchor.

Module scores and differential-expression counts are preprocessed differently, by necessity. Every module score in this paper is computed on the unfiltered gene universe; every differential-expression count, on the detection-filtered matrix. A rank-based score tolerates genes tied at zero. An empirical-Bayes variance model does not (§2.4). The two quantities are not two views of one matrix, and a reader comparing a background size in Supplementary Table S10 against a differential-expression count in the same row should expect them to differ. All percentages in the paper share a single scale.

One modality dataset could not be re-derived and should be discounted. On re-running the modality pipeline, the record for GSE262146 (listed as fluid shear in THP-1 cells, 3 versus 3) could not be reconciled with that accession: it deposits single-nucleus RNA and ATAC data from a different experimental system, and no companion bulk series was located. Its original contribution was a −1.5% interferon shift at permutation p = 0.90. No conclusion in this paper depended on it. It is now excluded from the modality projection of §3.5 and from the two-arm analysis of §3.9, leaving 22 contrasts from 20 datasets. Every other modality contrast was re-derived from its deposited matrix under the single scoring convention of §2.4. Three loaders required correction in the process: a spreadsheet path, a column-name pattern that no longer matched the deposited header, and an unmapped gene symbol admitted as a row name once the detection filter was removed.

What we did not do. No cross-study or cross-modality batch correction was applied anywhere: the rank-based single-sample score makes it unnecessary, and this reflects a design choice. Model-assumption diagnostics were not run, since the inference is permutation-based.

### 4.10 What would change this conclusion

The central claim is an equivalence, not a discovery, so it’s worth stating what observations would overturn it, in descending order of how likely we think each is.

A gradient design with adequate replication. The pooled estimate is a statement about two-level chords. A stiffness or force gradient with four or more levels and six or more biological replicates, including a chemical anchor in the same batch, would test the +11.4%-per-decade slope seen in the one four-point dataset available (§3.8). If that slope is real, the chord across a physiological range would still be small, which is why it survives the present analysis. But the mechanism would be quite different from “mechanics does not touch this arm.” This is the most valuable single experiment the field could run, and it is not expensive.

Four to six additional human studies would be needed. The human stratum carries K = 3 and returns an equivalence margin of ±20% rather than ±10%. A study-level permutation over three studies cannot return a p below 0.25: this is a resolution limit, not a finding. Adding human studies could move the human estimate outside the margin even if the mouse estimate does not move at all. The two species are not interchangeable here, and the mouse stratum is both tighter (I² = 12.0%) and larger.

A time course. Every contrast in this corpus is a single time point, and the interferon arm is kinetically sharp: in the chemical anchors the module is already near maximum by 4–6 h. A mechanical input that produces a transient ISG pulse resolving before the sampled time point would be invisible to every dataset analysed here, and no feature of our design would reveal it. We cannot bound this possibility.

A sorted-cell in-vivo measurement. The two largest absolute shifts in the modality projection, +55.9% and +38.8% of the anchor, both come from tissue preparations in which myeloid composition and macrophage-intrinsic response are confounded. Repeating either in sorted macrophages, with the same mechanical contrast, would convert the largest apparent effects in the corpus into interpretable ones, in whichever direction they then fall.

What would *not* change the conclusion is a further two-level stiffness experiment with effector-arm readouts, however carefully executed. Thirty-three such contrasts are already scored here: eleven stiffness and applied-force contrasts in §3.9 and twenty-two modality contrasts in §3.5. The design question they answer is not the one in dispute.

### 4.11 What this implies for experimental design

Three recommendations follow directly, and all three are cheap.

1. Include a chemical anchor in the same batch. Almost none of the datasets analysed here contains an LPS or LPS + IFN-γ arm processed alongside the mechanical arm. Without one, “mechanically induced M1” cannot be placed on any scale: a reader cannot tell a 5% effect from a 100% effect.
2. Report mechanical dose, and use more than two levels. Section 3.8 explains why: two-level designs cannot distinguish “no effect” from “wrong part of the curve.”
3. Do not use interferon-stimulated genes as the readout for a mechanical perturbation, and report both arms. ISGs are the part of M1 least responsive to mechanics in this corpus, so a null result there says nothing about whether the perturbation worked. The NF-κB/AP-1 effector arm is the one that responds, in either direction. Scoring both arms on the same samples costs nothing beyond the gene lists given in Supplementary Table S2. A paper that reports a single effector marker and calls the result “M1” cannot distinguish the two situations this work separates.
4. Deposit a myeloid-resolved measurement when the perturbation is applied in vivo. The two largest apparent effects in this corpus come from whole-tissue or whole-organ preparations, in which a shift in module score cannot be separated from a shift in composition; both studies were demoted to the sensitivity set for this reason. Sorting, or depositing single-cell data that permits myeloid pseudobulk, converts an uninterpretable measurement into an interpretable one. It costs nothing extra to do.

A fifth recommendation is addressed to the field rather than to an individual experiment. The quantity that made this analysis possible, a mechanical effect expressed as a fraction of a chemical effect measured the same way, required us to assemble anchors from studies that had nothing to do with mechanics. The mechanical studies themselves did not contain them. If the stiffness literature adopted a single shared reference contrast, processed in-batch, the next synthesis of this kind would need no anchor-reconstruction step. Individual papers would become directly comparable to one another, not only to a meta-analysis.

## Data Availability Statement

This study analysed publicly available datasets only; no new primary data were generated. All accessions analysed are listed in Supplementary Table S1 (search strings and per-query hit counts) and Supplementary Table S3 (the 58 datasets verified record by record, with eligibility decisions). Two datasets were quantified from raw reads by the authors: E-MTAB-7900 (ArrayExpress; FASTQ obtained from ENA) and GSE299513 (GEO; author-deposited transcript abundances aggregated to gene level). The human effector anchors are GSE112372 and GSE110947; the single-cell modality contrasts were pseudobulked by the authors from GSE275018, GSE274828 and GSE186185. All derived study-level effect sizes, leave-one-out refits, species strata, power curves, module-level results and the complete modality-projection scoring table are provided as Supplementary Data. Analysis code and all intermediate result tables are openly available at https://github.com/congzhaoqing/mechanical-vs-chemical-macrophage-m1, released under the MIT Licence for the code and CC BY 4.0 for the result tables. The repository covers the scoring library, the per-dataset loaders, the contrast specification keyed to sample accessions, the single-cell pseudobulk scripts, and the diagnostic scripts used for the reproduction audit.

## Ethics Statement

Ethical approval was not required for this study: it is a secondary analysis of de-identified, publicly archived transcriptomic datasets. The original studies obtained the approvals reported in their respective publications.

## Author Contributions

CZ: methodology, formal analysis, writing — original draft, and writing — review and editing. WC: validation, investigation, and writing — review and editing. TZ: investigation. SZ: writing — review and editing. JZ: data curation. ZC: conceptualization, methodology, software, visualization, supervision, and funding acquisition. The authors read and approved the submitted version.

## Acknowledgments

Generative AI statement. The authors used Claude (Anthropic), accessed through the Anthropic API and a Claude-based agentic analysis environment, during the preparation of this work. Its use covered writing and debugging the R and Python analysis code described in §2, assembling and checking the dataset inventory. All gene sets, contrasts, datasets and statistical choices were specified by the authors; every numerical result reported here was produced by the deposited code and checked by the authors against the supplementary tables.

## Supplementary Material

The Supplementary Material for this article comprises Supplementary Tables S1-S15. S1 gives the 48 pre-specified search strings with per-query hit counts; S2 the curated gene-set definitions with source DOI and supplementary-table location; S3 the 58 datasets verified record by record with their eligibility decisions; S4-S8 the study-level effect sizes, leave-one-study-out refits, species strata, module-level results with small-study-effect tests, and the Monte-Carlo power curve for the main pool; S9 the chemical reference anchors; S10 the complete modality-projection scoring table; S11 the study-level effect sizes for all four modules in the expanded K = 12 pool; S12 the effector-ruler validation across the IFN-γ and LPS anchor cohorts; S13 the study-level two-arm effects for the eleven stiffness and applied-force contrasts; S14 the two-arm projection across all 21 modality contrasts with each design’s attainable minimum p; and S15 the five-module result for GSE274336, the corpus’s most transcriptionally responsive contrast, with its detected gene background. No supplementary figures are provided: the robustness analysis that earlier circulated as a supplementary figure is Figure 6 of the main text.

